# Regional heterogeneity shapes macroscopic wave dynamics of the human and non-human primate cortex

**DOI:** 10.64898/2026.01.22.701178

**Authors:** Victor Barnes, Jace Cruddas, Trang Cao, Isaac Z. Pope, Ting Xu, Thomas Funck, Nicola Palomero-Gallagher, James C. Pang, Alex Fornito

**Affiliations:** School of Psychological Sciences, The Turner Institute for Brain and Mental Health, and Monash Biomedical Imaging, Monash University, Clayton, Victoria, Australia; Center for Integrative Developing Brain, Child Mind Institute, New York, NY, USA; Institute of Neuroscience and Medicine (INM-1), Research Centre Jülich, Jülich, Germany; C. & O. Vogt Institute of Brain Research, Heinrich-Heine-University Düsseldorf, Düsseldorf, Germany

**Author notes:** Equal author contributions (James C. Pang, Alex Fornito).

## Abstract

Growing evidence indicates that macroscopic cortical activity is dominated by propagating waves of excitation. However, many computational models of such wave dynamics assume that the cortex is a spatially homogeneous medium, ignoring the rich regional variations in cellular, molecular, and physiological properties that are known to shape how brain activity evolves through both space and time. Here, we develop a general framework grounded in neural field theory to model how regional heterogeneities in diverse cortical properties shape spatiotemporal brain activity evolving on cortical surface meshes. This enables high resolution, vertex-level simulations without requiring predefined parcellations. The model requires only a standard mesh model of the cortical manifold and a spatial heterogeneity map, providing a biologically grounded and computationally efficient framework that can be generalised to human and non-human species. Using multiple cellular, molecular, and physiological maps in humans—and analogous maps in macaques and marmosets—we show that our model can consistently recapitulate known relationships between regional heterogeneities and variations in cortical wave speed. In particular, we find that models parameterised by heterogeneities in intracortical myelin and excitation-inhibition balance yield the largest performance improvements relative to spatially homogeneous models. Our results identify a key role for regional variations in myelin, receptor, and genetic architecture in shaping the spatial patterning of macroscale cortex-wide activity that is conserved across primate species with diverse cortical geometries.

## Introduction

Over a century ago, Brodmann published his seminal work parcellating the cerebral cortex of different mammals into discrete areas according to regional variations in cytoarchitecture^1^. He viewed these areas as representing “specific morphological organs” that each possess a distinct function (p. 251; ref.^1^). Although such a mapping between structure and function is now considered too simplistic^2-7^, understanding how regional specialisation of cortical function is related to spatial variations in cellular, molecular, and genetic properties has been a fundamental focus of subsequent neuroscientific research. In this context, the growing availability of diverse types of brain maps acquired in different species has revealed that distinct molecular, cellular, genetic, physiological, and metabolic properties vary spatially across the cortex in characteristic ways that correspond with various measures of brain activity^3,8-11^.

Large-scale biophysical models of cortical dynamics offer a formal way of interrogating the influence of heterogeneous cortical properties on regional neural activity^12^. Neural mass models (NMMs) represent one prominent class of such models. These models divide the cortex into discrete areas with intrinsic dynamics governed by a set of biophysical equations describing their mean-field properties (e.g., membrane potential, firing rate). Neuronal populations within an area interact with those in other areas via their axonal inter-connectivity, as measured with invasive tract tracing^13^ or non-invasive diffusion magnetic resonance imaging (MRI)^12,14-16^.

Initial applications of NMMs treated all cortical areas as having homogeneous internal structure, with heterogeneity driven purely by variations in inter-regional connectivity^17-19^ but recent work has extended NMMs to explore how spatial variations in distinct cortical properties may influence region-specific biophysical processes and facilitate better prediction of empirical measures of brain activity. For instance, Demirtaş et al.^20^ showed that incorporating hierarchical heterogeneity in regional synaptic strengths improves predictions of empirical inter-regional functional coupling (FC) estimates measured in humans with resting-state functional MRI (fMRI), while also enhancing predictions of spectral features observed in resting-state MEG. The synaptic strengths were parameterised using the ratio of T1-weighted (T1w) MRI and T2-weighted (T2w) MRI, which has been proposed as a non-invasive index of intracortical myelin^21^ and hierarchical rank within cortical processing hierarchies^22^. Similarly, Deco et al.^23^ demonstrated that transcriptional heterogeneity in the ratio of gene expression markers for excitatory and inhibitory receptors, used as a proxy for regional excitation:inhibition (E:I) balance, allows NMMs to more accurately reproduce diverse spatial and temporal properties of human resting-state FC. Wang et al.^24^ developed an inversion approach to estimate regional variations in multiple biophysical parameters of NMMs that best fit resting-state FC, and found that the optimal spatial pattern of regional variations in recurrent excitation strength closely tracks the classical cortical hierarchy. A similar approach by Saberi et al.^25^ showed that regional heterogeneity in the E:I balance of input current within sensory and association areas can capture patterns of neurodevelopmental maturation in adolescence. Other work has shown that incorporating regional heterogeneities in NMMs can overcome some limitations of NMMs, such as over-fitting of widespread signal deflections in fMRI data^26^, and may also explain phase transitions in neural synchronisation^27^.

One limitation of NMMs is that they are generally only computationally feasible for networks comprising a few hundred regions, meaning that they often rely on a coarse areal discretisation of the cortex that can miss the effect of fine-scale variations in the underlying map of cortical regional heterogeneity. The reliance on a discrete areal parcellation creates further challenges, as the reliability and utility of such parcellations has been debated since the time of Brodmann, and there is a growing list of differing atlases available in the literature, making the choice of a specific parcellation somewhat arbitrary (for overviews see refs.^3,7,11,28-30^). Indeed, wide-field recordings of mesoscopic and macroscopic dynamics indicate that activity often propagates as travelling waves of excitation that can traverse areal boundaries^31,32^, and theoretical work shows that waves naturally emerge even within discretised NMM dynamics^33^, suggesting that the imposition of discrete boundaries on the cortex may limit our ability to accurately model how the evolution of cortical activity through both time *and* space is shaped by fine-scale spatial variations in cellular, molecular, and physiological properties^32^.

Neural field theory (NFT)^34-36^ refers to a class of models that overcome these limitations by eschewing discrete parcellations and treating the cortex as a continuous sheet through which travelling waves of activity propagate^37,38^. A particular variant of NFT by Robinson and colleagues^36^ has been widely applied to electroencephalography (EEG) and local field potentials (LFPs) to successfully model diverse phenomena such as the emergence of canonical EEG rhythms^37^, evoked potentials^39,40^, arousal states^41,42^, and waves of cortical activity^43^. Traditional NFT-based models were formulated on simple geometries such as 2D planes and spheres, but recent advances have extended their application to the complex geometry of the cortical manifold^44-47^. Such geometry-aware NFT-based models have been shown to more parsimoniously explain diverse aspects of cortical activity measured with fMRI than popular and more highly parameterised NMMs^46^.

The majority of NFT models applied to data thus far have assumed that the cortex is a homogeneous sheet, ignoring spatial variations in the cellular, molecular, and microstructural properties that NMM studies have shown strongly influence cortical dynamics^20,23,24^. Here, we address this limitation by deriving a simple, two-parameter NFT-based model that can account for spatial heterogeneities in any arbitrary cortical property. The simplicity of our model means that it can simulate activity at high spatial resolution (i.e., at resolutions finer than the scale of discrete cortical areas), affording a fine-grained characterisation of how regional variations in different cortical properties influence dynamics. Importantly, our framework generalises across species, enabling systematic tests of whether conserved organisational features of regional heterogeneity shape large-scale dynamics across species. By applying the same model to humans, macaques, and marmosets, we aim to identify shared microstructural and molecular axes that constrain cortical activity across diverse cortical geometries. To this end, we compare the effects of eight heterogeneity maps defined for the human cortex, spanning myeloarchitecture, cytoarchitecture, chemoarchitecture, microstructure, macrostructure, and dynamics; nineteen maps for the macaque cortex, including structural and receptor-based properties; and three maps for the marmoset cortex, including structural and cytoarchitectural properties. We show that accounting for such heterogeneity generally allows one to capture fMRI data more accurately than spatially homogeneous models, and that models in which the local speed of wave propagation is influenced by either intracortical myelin content (as quantified using the T1w/T2w ratio) or markers of regional E:I balance yield the greatest improvement in model performance. Critically, we provide evidence that these effects are consistent across primates, suggesting that the principles that underlie how regional heterogeneity shapes large-scale dynamics are conserved across primate species separated by over 47 million years of evolution.

## Results

### Theory

We model mean-field macroscale brain activity as waves propagating through the cortical sheet, with the propagation described by a damped wave equation derived from Robinson et al.’s NFT formulation^37,38,46^. Here, we extend the NFT wave model by generalizing the form of the damped wave equation to incorporate the effects of spatially varying local properties on the speed of wave propagation, defined as

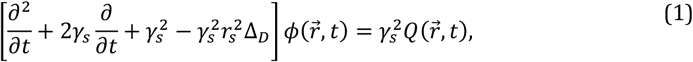

where 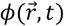 is the neural activity at location 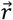 and time 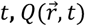 is the external input, *γ*_*s*_is the damping rate, *r*_*s*_ determines the spatial length scale of the assumed connectivity kernel between different cortical points (and thus, the wavelength of the wave propagation), *γ*_*s*_*r*_*s*_ is the average wave speed of the modelled dynamics, and 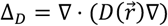 is the generalised Laplace-Beltrami Operator (LBO; see Methods). The LBO describes geometric variations of the cortical manifold, which play an important role in shaping wave dynamics^46^. The diffusion tensor 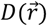 generalises the LBO so that it can account for local heterogeneities in the cortical sheet, as defined by a given spatial map encoding variations in some property of interest (e.g., myeloarchitecture or the expression levels of a particular gene).

We define the diffusion tensor as a diagonal matrix whose elements follow a sigmoidal transform, 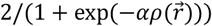, where *α* is a free parameter that scales the given z-scored cortical map 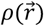. This formulation allows us to flexibly modulate the relationship between local heterogeneity and wave speed, such that positive *α* values increase wave speed in regions of large 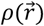, while negative *α* values invert the relationship. As *α* increases in magnitude, the resulting distribution of the values of the diffusion tensor becomes more skewed, accentuating regional differences in propagation speed (see Methods; Fig. S1). Note that in the special case of *α* = 0, 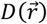 becomes the identity matrix, such that the generalised LBO converges to the standard LBO^46^, Δ, and our model reduces to the homogeneous wave equation, as evaluated in previous work^46^.

We simulate the NFT wave model on a group-average representation of the human cortical midthickness surface of the left hemisphere derived from T1-weighted MRI data and discretised into a triangular mesh comprising 4002 vertices (fsLR-4k) using established procedures^48,49^ (Fig. 1a). The resulting neural activity is then converted to simulated blood-oxygenation-level-dependent (BOLD) signals using the popular Balloon-Windkessel hemodynamic model^50,51^ to make it directly comparable to empirical resting-state fMRI data (Fig. 1a). A toy example of how wave propagation can be affected by spatial heterogeneities in the cortical medium is presented in Fig. 1b. In the homogeneous case, waves triggered by an initial impulse propagate isotropically and with uniform speed along all directions in the medium. In the heterogeneous case, spatial variations disrupt this isotropy and uniformity, leading to local variations in wave speed and a distinct propagation front.

**Fig. 1.**
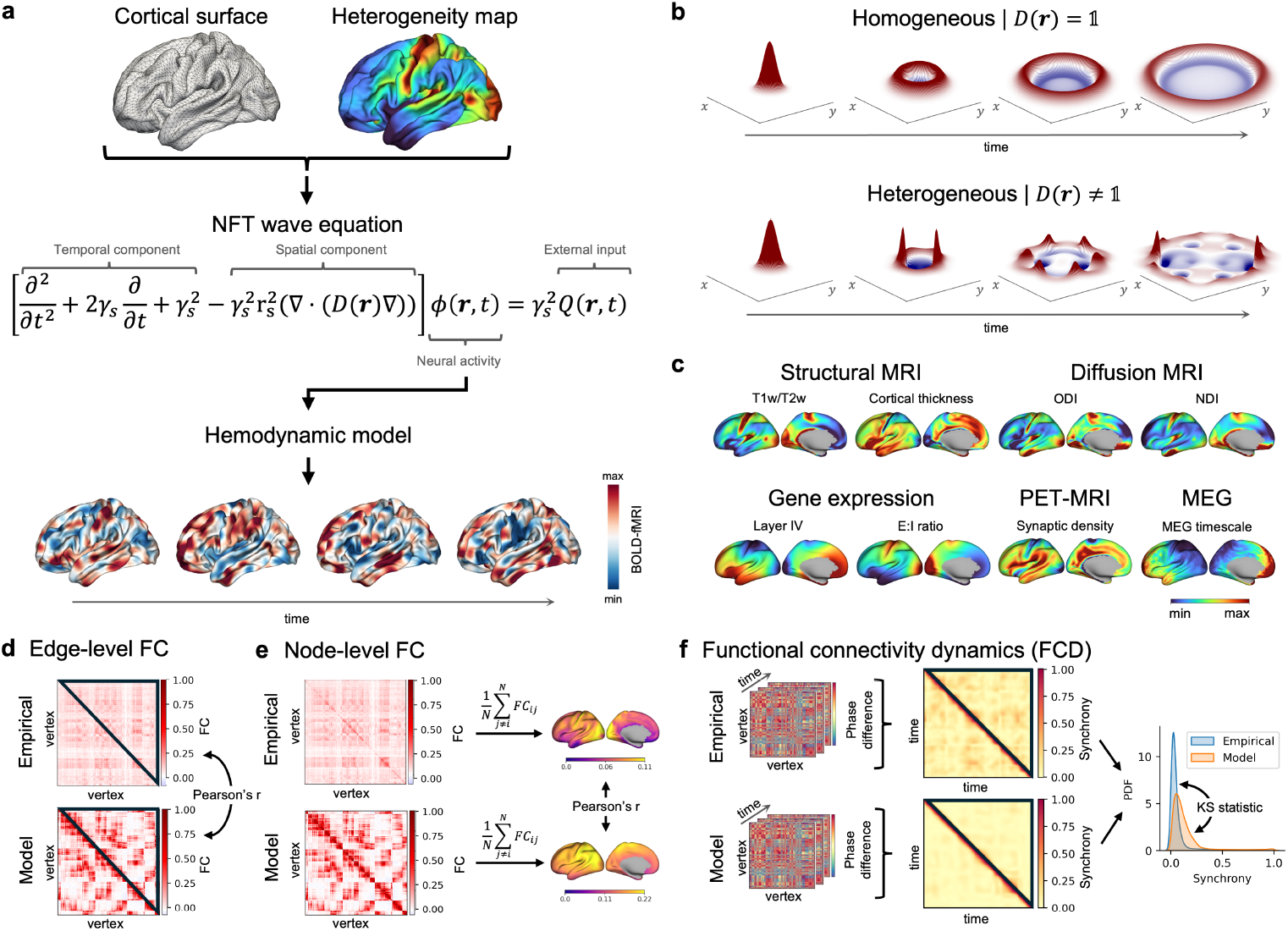
A spatially heterogeneous wave model of large-scale cortical activity. **a**, We use a generalised wave equation to simulate dynamics on a mesh model of the cortical surface under differing assumptions of either spatial homogeneity or different forms of spatial heterogeneity. The resulting dynamics are then passed through a hemodynamic model to simulate spatiotemporally evolving BOLD-fMRI signal dynamics. **b**, Toy example comparing wave propagation in the homogeneous case (when the diffusion tensor, 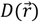, is equal to the identity matrix) and the heterogeneous case (when the diffusion tensor diverges from the identity matrix). Spatial heterogeneity modulates local wave speed, leading to distinct wave patterns. **c**, Heterogeneity maps describing different cortical properties hypothesised to have an impact on wave propagation. These maps are used to generate the eight heterogeneous models. Note that these maps are visualised on the inflated version of the fsLR-4k surface for clarity. **d-f**, Evaluation metrics used to fit each model to empirical resting-state fMRI data: **d**, Edge-level FC is calculated by computing the Pearson’s correlation, *r*, between the upper-triangular elements of the model and empirical FC matrices, **e**, Node-level FC is calculated by computing the Pearson’s correlation between the mean FC strength of each cortical vertex between the model and empirical data, **f**, Functional connectivity dynamics (FCD) are summarised by filtering vertex-level time series (0.04-0.07 Hz), extracting instantaneous phases via the Hilbert transform, and quantifying the similarity of pairwise phase synchrony over time, with model-data agreement assessed using the Kolmogorov-Smirnov (KS) statistic.

Note that our model does not require an a priori measurement of inter-regional connectivity (i.e., a connectome) as the spatial component of the dynamics is determined entirely by the geometry of the cortex and the heterogeneity map, as encoded in the generalised LBO. However, our approach does account for a simple connectivity kernel in which different cortical locations are coupled through an isotropic, distance-dependent kernel approximating the well-known exponential distance rule proposed as a dominant and universal organisation principle of connectomes^52,53^. Prior work has indicated that this approximation is sufficient to capture fMRI dynamics with accuracy comparable or superior to NMMs that rely on direct measurement of the connectome^46^. The advantage of our geometric approximation is that we only require a model of cortical geometry as input, which can easily be obtained from T1-weighted MRI, thus greatly enhancing its applicability to non-human species (e.g., macaque and marmoset to be analysed in the next sections) for which connectomes have only been mapped with partial cortical coverage and at coarse, area-level resolutions^54,55^.

We incorporate spatial heterogeneity into the NFT wave model using one of eight different maps, encoding cortical variations in distinct molecular, cellular, microstructural, anatomical, and physiological properties in the human brain (see Fig. 1c for each spatial map and Methods for more details on their derivation), which we hypothesise to have the greatest potential effects on wave propagation. Specifically, these maps are: (i) the T1w/T2w ratio derived from MRI, which serves as an approximation of intracortical myelin content^56^, and is thus putatively related to axonal conduction velocity; (ii) cortical thickness, measured with T1w MRI, which reflects regional microstructural organization, including neuronal density and packing^56^; (iii) synaptic density, measured with [^11^C]UCB-J positron emission tomography (PET) of the Synaptic Vesicle glycoprotein 2A (SV2A)^57^, which approximates regional capacity for local information transfer; (iv) E:I balance quantified using the expression ratio of gene markers for excitatory and inhibitory neurons^58^, which approximates regional variations in local excitability/damping; (v) expression of Layer IV-specific genes^58^, which approximates regional variations in thalamic input; (vi) orientation dispersion index (ODI), derived from diffusion MRI^59^, which indexes the heterogeneity of neurite fibre orientations; (vii) neurite density index (NDI) derived from diffusion MRI^59^, which indexes the proportion of the voxel volume occupied by neurites; and (viii) regional intrinsic activity timescale derived from magnetoencephalography (MEG), which measures local dynamic properties^60^.

For each model, we simulate resting-state activity driven by a stochastic input modelled as white Gaussian noise, following previous work^46,61-63^, and compare the simulations to empirical fMRI data acquired in 255 unrelated participants from the Human Connectome Project (HCP)^64^. We use the aforementioned group-averaged cortical surface as the spatial manifold for the dynamics. Our model contains two free parameters: *α* and *r*_*s*_, which are optimised to fit the group-average fMRI data, with the objective function defined as a summation of two measures of static FC—edge-level FC fit (*r*_*edge*_; Fig. 1d) and node-level FC fit (*r*_*node*_; Fig. 1e), sometimes referred to as FC strength^65,66^ or global brain connectivity^20,67^—and one time-resolved measure of FC—functional connectivity dynamics (FCD) fit (*KS*_*FCD*_, Fig. 1f; see Methods for further details). These three metrics are commonly used in the literature to test the accuracy of large-scale brain models of cortical dynamics^23,25,26,46^. To avoid overfitting and allow fair comparison between homogeneous and heterogeneous models, which differ in their number of free parameters, we employ a 5-fold cross-validation and use 80% of participants for training and 20% for testing in each fold. Our workflow enables a systematic comparison of how various local cortical properties influence macroscopic wave propagation and their ability to improve model-based predictions of observed brain activity patterns.

### Shared and unique aspects of cortical heterogeneity

We first quantify the spatial similarity between the different heterogeneity maps by computing absolute pairwise Pearson’s correlations between maps. To better visualise maps that have highly comparable structures, we reorder the rows and columns of the resulting correlation matrix using hierarchical clustering, as shown in Fig. 2a. We use absolute correlations here since our model can learn either a direct or inverse relationship between the heterogeneity map and wave speed, depending on the sign of *α*. Figure 2a shows that cortical thickness, E:I ratio, T1w/T2w, Layer IV, and MEG timescale maps form one cluster characterised by moderate-to-strong inter-map correlations (i.e., 0.42 < r < 0.79). The other maps are weakly correlated. The exceptions are the two microstructural maps derived from diffusion MRI—i.e., ODI and NDI— which show a spatial correlation of r=0.70. This analysis indicates that our maps capture both shared and unique sources of regional heterogeneity in human cortex.

**Fig. 2.**
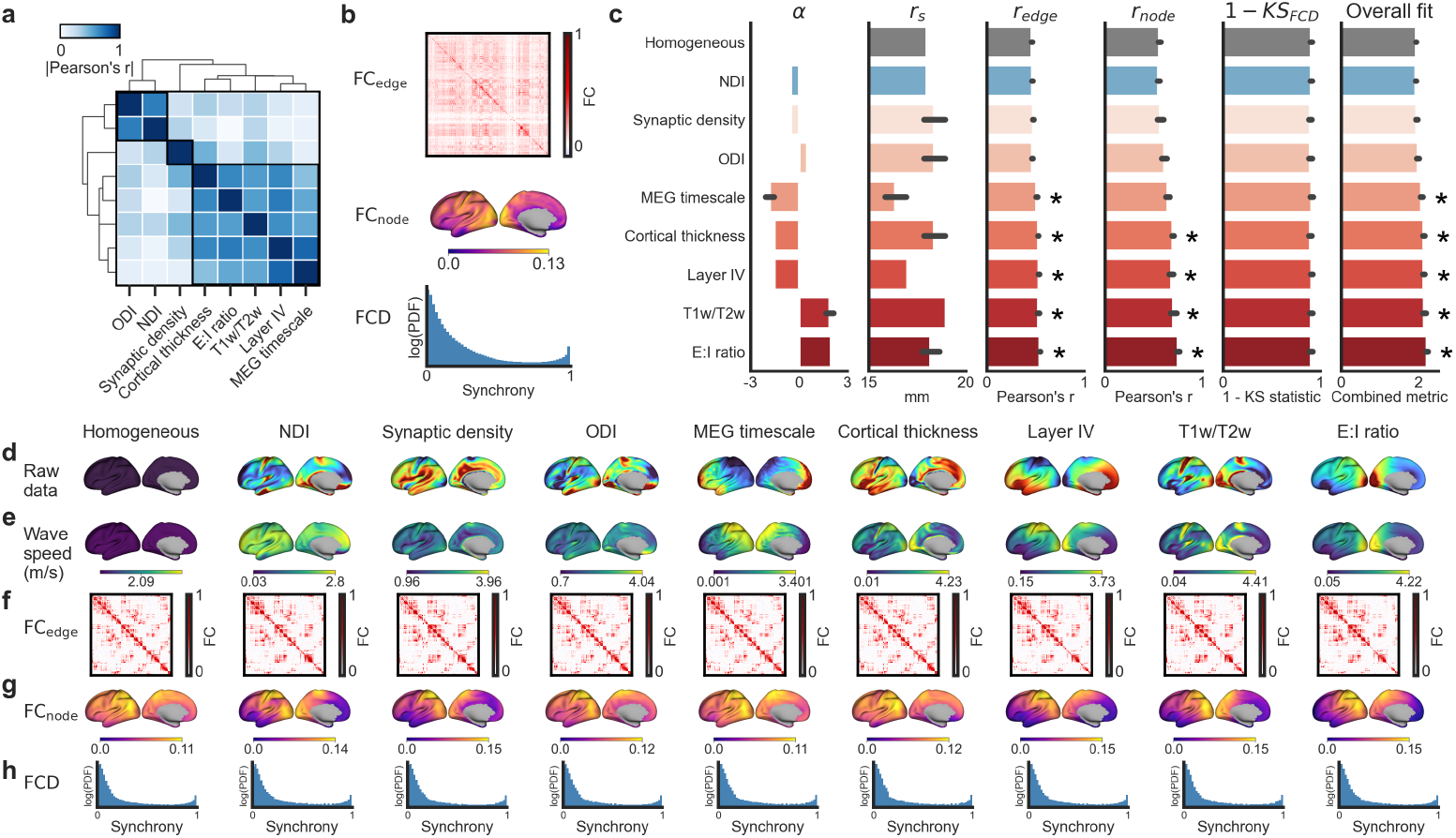
Incorporating regional heterogeneity captures spatially resolved dynamics more accurately. **a**, Inter-map similarity (absolute Pearson’s correlation) across cortical heterogeneity maps ordered by hierarchical clustering, revealing a dominant cluster of maps with moderate-to-strong correlations along with a small cluster of diffusion MRI-derived measures of microstructure. **b**, Empirical vertex-level target properties used for model fitting and evaluation: (top) group-averaged edge-level functional coupling (FC) matrix (*FC*_*edge*_), (middle) vertex-wise node-level FC (*FC*_*node*_), and (bottom) distribution of functional connectivity dynamics (FCD; log-scaled y-axis). **c**, Optimal parameters (*α* and *r*_*s*_) and cross-validated fits for each model (bars: mean across 5 folds; error bars: s.d.). The homogeneous model is shown for reference (grey), followed by the heterogeneous models ordered from lowest to highest overall fit (blue: lower overall fit than homogeneous; red: higher overall fit than homogeneous). The FCD fit is shown as 1 - KS statistic since lower KS values indicate a better fit. The overall fit is calculated as ℒ = *r*_*edge*_ + *r*_*node*_ + (1 − *KS*_*FCD*_). For each evaluation metric, models that significantly outperformed the homogeneous model, assessed via eigenstrapping nulls (see Methods), are marked by an asterisk (p < 0.05) **d-h**, Model inputs and evaluation outputs for the homogeneous and heterogeneous models. **d**, Raw heterogeneity maps displayed on the cortical surface. **e**, Resulting spatial maps of wave speeds derived by using the optimised alpha parameter (see Methods). **f-h**, Model-derived (f) edge-level FC matrices, (g) node-level FC maps, and (h) FCD distributions (log-scaled y-axis) all defined at the vertex level. Panels d-h are arranged from left to right following the ordering in panel c.

### Mapping cortical heterogeneity to wave dynamics

We fit each model parameterising heterogeneity using one of the eight maps (Fig 1c) to the group-average edge-level FC, node-level FC, and FCD, calculated across the concatenated timeseries from each subject (Fig. 2b; see Methods). We present the full optimisation landscapes for *α* and *r*_*s*_in Fig. S2, which show how each fit metric varies across the parameter space and that the selected optima lie within well-defined minima (see Methods).

Across models, the fitted *α* parameter determines how each cortical property influences local propagation speed: positive *α* values indicate a direct relationship, such that regions with higher map values support faster wave propagation; negative *α* values indicate an inverse relationship, in which higher map values lead to slower wave propagation. The relationship between intracortical myelin content, as indexed by the T1w/T2w map, and wave speed is well-known^68^, where areas with high T1w/T2w signal are expected to yield faster wave propagation. As expected, our optimisation procedure yields a positive *α* for this map (Fig. 2c). Similarly, we identify a positive *α* for the E:I ratio map, which aligns with the expectation that higher local excitability should facilitate wave propagation (Fig. 2c). These results indicate that our optimisation procedure captures biologically plausible relationships between cortical heterogeneity and wave dynamics.

The expected relationships between wave speed and other cortical heterogeneity maps are less clear. Our optimisation identifies a positive *α* for the ODI maps (Fig. 2c), but negative *α* for cortical thickness, NDI, synaptic density, layer IV cell density, and MEG timescale (Fig. 2c), indicating that higher values of these properties correspond to slower waves in the fitted dynamics. These mappings can be seen when comparing the raw heterogeneity maps (Fig. 2d) with their corresponding wave speed fields (Fig. 2e): models with *α* > 0 produce wave speed maps that track the raw heterogeneity, whereas *α* < 0 models produce an inverted pattern.

The parameter *r*_*s*_ determines the length scale of connectivity in the model. Notably, the MEG timescale and Layer IV models exhibit lower optimised *r*_*s*_values compared to the homogeneous model (Fig. 2c). This result suggests that the MEG and Layer IV models rely on more localised interactions to reproduce the empirical fMRI data. Because *r*_*s*_ scales the mean propagation speed (via *γ*_*s*_*r*_*s*_), optimising *r*_*s*_ effectively shifts the average wave speed of the modelled dynamics. Notably, the fitted values yield mean wave speeds that fall within physiologically plausible ranges and are consistent with empirical measurements and prior modelling work^68-70^ (Fig. 2e).

### Spatially heterogeneous models outperform homogeneous models at high spatial resolution

We first evaluate the performance of the homogeneous wave model in capturing diverse aspects of resting-state fMRI dynamics as a benchmark for the heterogeneous models. The homogeneous model shows edge-level FC and node-level FC correlations of 0.434 ± 0.004 and 0.562 ± 0.013, respectively, along with a KS statistic of 0.089 ± 0.009 for FCD, which is consistent with past estimates of the performance of this model^46^ (see Fig. 2f, g, and h for the model edge-level FC, node-level FC, and FCD results, respectively, as well as Fig. 2b for the corresponding empirical results).

Introducing spatial heterogeneity improves overall model fits relative to the homogeneous case for most heterogeneity maps (Fig. 2c). Using the aggregate measure of fit defined as our objective function, the best performing model is parametrised by the E:I ratio map derived from transcriptomic data (see Methods), consistent with past work on NMMs^23^ (Fig. 2c). This model achieves edge-level FC and node-level FC correlations of *r*_*edge*_ = 0.547 ± 0.003 and *r*_*node*_ = 0.757 ± 0.013, representing improvements of 26.0% and 34.7% over the homogeneous model, respectively. Models parametrised by T1w/T2w (*r*_*edge*_ = 0.531 ± 0.009, *r*_*node*_ = 0.707 ± 0.034), layer IV (*r*_*edge*_ = 0.536 ± 0.006, *r*_*node*_ = 0.684 ± 0.025), cortical thickness (*r*_*edge*_ = 0.529 ± 0.004, *r*_*node*_ = 0.700 ± 0.012), and MEG timescale (*r*_*edge*_ = 0.514 ± 0.006, *r*_*node*_ = 0.648 ± 0.023) closely follow, suggesting that each of these maps yields similar performance. Indeed, these five best performing maps are spatially correlated with each other (Fig. 2b), and all align with a unimodal-transmodal gradient (Fig. 2b). This gradient corresponds to a principal axis of organisation for many different cortical properties and is thought to support hierarchical processing of sensory information^3,22,71^. The improvements relative to the homogeneous model associated with the NDI, ODI, and synaptic density maps are marginal (Fig. 2c), with their performance being particularly poor in capturing node-averaged FC maps relative to the other heterogeneous models (Fig. 2c). This is not to say that spatial variations of these properties have no effect on dynamics; rather, these findings indicate that such effects are not readily captured by regional changes in wave speed, as parameterised by our model.

Parsing the objective function into its constituent features shows that the performance gains of the E:I and other similarly performing models primarily arise through their improved ability to capture static rather than dynamic properties of FC (Fig. 2c). This specificity likely results from the fact that edge-level and node-level FC capture information about network topography (i.e., how FC is spatially distributed throughout the brain), which will be more strongly influenced by spatial heterogeneity in the maps used to parametrise the model. In particular, the largest improvements in model fit appear in node-level FC.

In contrast to static topographic properties, FCD captures statistical properties of the dynamics but does not explicitly incorporate a measure of spatial similarity and should thus be less affected by variations in the spatial properties of the model. Due to this limitation in capturing differences in spatiotemporal patterns, the FCD score remains largely invariant across models. Notably, the homogeneous model already achieves a low KS statistic (high “1 - KS statistic” in Fig. 2c), leaving little room for improvement when heterogeneity is added. Our findings thus indicate that incorporating spatial heterogeneity into the wave model improves the topographic properties of network activity while leaving the temporal statistics largely unchanged (Fig. 2c,h).

It is important to note that there is an inherent trade-off in individual metric performance because our optimisation procedure jointly maximises an aggregate objective across three metrics (edge-level FC fit, node-level FC fit, and FCD fit). This multi-objective approach ensures balanced performance across static and dynamic properties but prevents the model from achieving the absolute best fit for any single metric. Optimising exclusively for edge-level FC would yield substantially higher correlations for that measure, while prioritising node-level FC would similarly improve those respective fits. For instance, optimising the E:I model on each of the metrics individually yields fits of *r*_*edge*_ = 0.575 ± 0.011 and *r*_*node*_ = 0.772 ± 0.011; whereas doing the same for the T1w/T2w model yields fits of *r*_*edge*_ = 0.557 ± 0.014 and *r*_*node*_ = 0.736 ± 0.027. Our decision to optimise across all three metrics follows past work^23,46^ reflects a deliberate compromise aimed at capturing complementary aspects of functional dynamics within a unified framework.

To ensure that the performance of our heterogeneous models is not driven by generic properties associated with the use of spatially autocorrelated maps, we use eigenstrapping^72^ to generate an ensemble of 500 randomised maps with the same spatial autocorrelation as the empirical heterogeneity maps (Fig. S3; see Methods). Each null map was used to parameterise a unique null model and was run through the same 5-fold cross-validation procedure, thus providing a benchmark for establishing the performance gains expected from using any generic autocorrelated heterogeneity map. This analysis reveals that, for edge-level and node-level FC fit, the E:I, T1w/T2w, layer IV, cortical thickness, and MEG timescale models all significantly outperform the homogeneous model. These five maps correspond precisely to those forming the cluster of moderate-to-strong correlations in the inter-map analysis (Fig. 2a). In contrast, we found no significant improvement in FCD fit across any of the heterogeneous models.

### Transmodal systems benefit most from regional heterogeneity

We next examine whether heterogeneous modelling improves fits to FC globally or preferentially within specific cortical systems by considering the edge-level FC similarity between model and data within canonical resting-state networks (RSNs; Fig. 3a). RSN assignments are based on a community detection solution derived from FC estimates in the HCP dataset using the HCP-MMP1 parcellation^20,56^. We focus on comparisons between the homogeneous model and the best performing heterogeneous (E:I ratio) model. For each vertex, we compute the correlation between its empirical and model-generated FC profiles, yielding a spatial map of vertex-level model accuracy (Fig. 3b; see Methods). We then take the difference in model accuracy between the heterogenous and homogeneous models to estimate model improvement, with positive values meaning that the heterogeneous model outperforms the homogeneous model (Fig. 3c). Qualitatively, we find that the E:I ratio improves fits across nearly the entire cortex, although association networks such as the frontoparietal, default mode, cingulo-opercular networks show more pronounced gains than sensory networks (visual, auditory, somatomotor). The E:I ratio map thus preferentially improves the model’s ability to capture the FC of higher-order cortical areas.

**Fig. 3.**
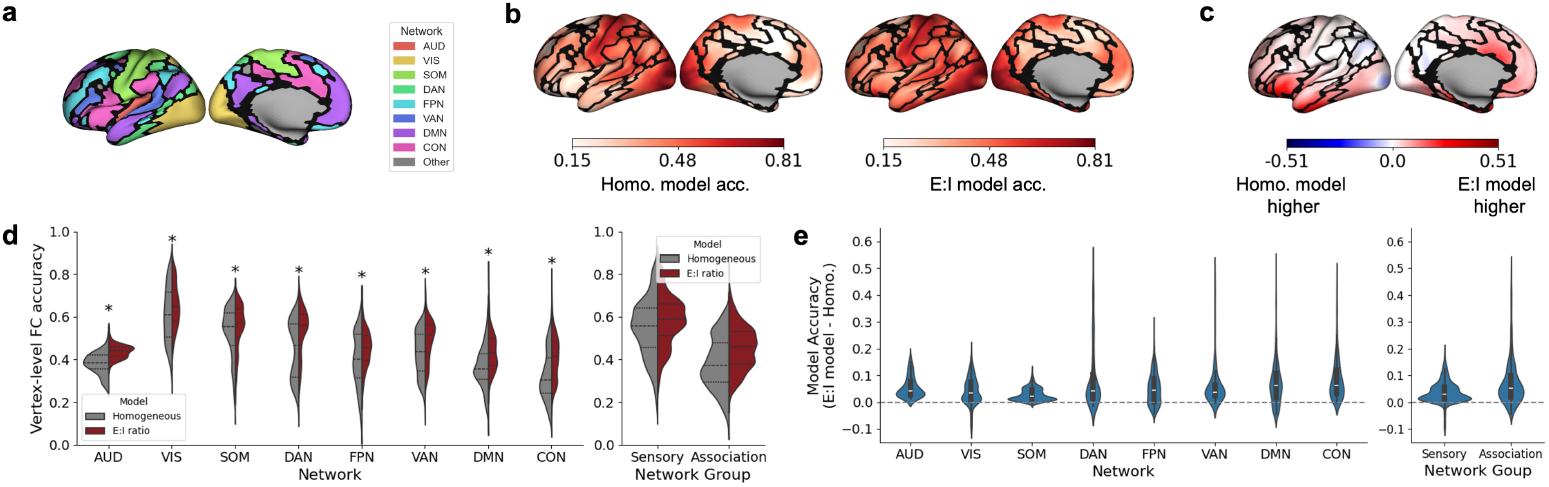
Accounting for local variations in E:I ratio preferentially improves model fits in the association cortex. **a**, Resting-state network (RSN) partitions used to summarise model performance, colour-coded by network: Auditory (AUD), Visual (VIS), Somatomotor (SOM), Dorsal Attention (DAN), Frontoparietal (FPN), Ventral Attention (VAN), Default Mode Network (DMN), and Cingulo-opercular (CON). **b**, Spatial distribution of vertex-level FC accuracy, defined as the model-empirical FC similarity, for the homogeneous and heterogeneous E:I models. **c**, Spatial distribution of vertex-level FC improvement, defined as the difference in model-empirical FC similarity between the best performing heterogeneous model (based on the E:I ratio map) and the homogeneous baseline. Positive values (red) indicate regions where the heterogeneous model better captures empirical FC patterns; negative values (blue) indicate regions where the homogeneous model performs better. **d**, Comparison of vertex-level FC accuracy at the individual network and network group levels (Sensory: AUD, VIS, SOM; Association: DAN, FPN, VAN, DMN, CON). Significant improvements in the heterogeneous model are marked by an asterisk (p < 0.001; two-sample t-test). **e**, Comparison of vertex-level FC improvements (E:I ratio - homogeneous) at the individual network and network group levels.

To quantify these effects, we aggregate vertex-level FC accuracy estimates within each RSN and compare the distributions between models (Fig. 3d, e). On average, sensory networks exhibit higher overall FC accuracy than association networks in both models (Fig. 3d). Despite these differences in baseline accuracy, all networks show significant improvements when using the heterogeneous model (Fig. 3d), with association networks exhibiting larger gains than sensory networks. Comparison of vertex-level improvements further confirms this trend (Fig. 3e), where their distribution reveals broader and more substantial gains in association networks.

### Advantages of vertex-level over parcel-level modelling

Our analysis thus far shows that our model can capture diverse FC properties at high spatial resolutions (i.e., >4,000 vertices) with an accuracy comparable to typical atlas-based analyses^20,23^. We now consider model performance in relation to traditional, atlas-derived FC estimates, which are more prevalent in the literature but can only model the dynamics of a few hundred regions. In particular, we consider how the choice of a specific atlas affects model fits. This question is important because many modelling approaches rely on predefined cortical parcels, which impose artificial boundaries that may not align with the underlying biological gradient of cortical heterogeneity.

To this end, we parcellate the data using the HCP-MMP1 and Schaefer atlases with 180 and 150 regions in one hemisphere, respectively, because they are two of the most popularly used in the field, they define an approximately similar number of regions, and they have distinct regional borders derived using different procedures and data modalities^11^. The parcellations are applied after each model run during each cross-validation fold such that each model is optimised on the parcellated edge-level FC, node-level FC, and FCD.

Figure 4a shows that the best performing heterogeneity map varies depending on the parcellation used. Specifically, the best performing model with the HCP-MMP1 atlas uses the Layer IV map and the best performing model with the Schaefer atlas uses the E:I ratio map. Notably, the HCP-MMP1 parcellation generally yields better model fits than the vertex-level model, despite the Schaefer atlas previously being shown to identify parcels with higher internal homogeneity, a key validation criterion for cortical atlases^11^. The superior performance observed with the HCP-MMP1 atlas likely reflects the fact that its parcel boundaries are defined using multimodal data coupled with manual adjustments designed to optimise known borders of primary sensory cytoarchitectonic fields. The borders thus track variations along the unimodal-to-transmodal gradient that aligns with the best performing heterogeneity maps in our models (Fig. 4b). In contrast, the Schaefer parcellation, which is based solely on data-driven clustering of vertex-level FC estimates, shows poorer alignment with the heterogeneity maps (Fig. 4c) and correspondingly has lower model performance than both the HCP-MMP1 atlas and our vertex-resolution model (Fig. 4a). These findings suggest that parcellations can either enhance or obscure biologically meaningful spatial variation depending on how well their boundaries match the underlying cortical features. Vertex-level modelling avoids this issue by preserving the continuous nature of cortical organisation, allowing us to more precisely capture the spatial heterogeneity with only a small cost in terms of model fit (relative to the HCP-MMP1). This analysis supports the use of vertex-level approaches when investigating the influence of spatially heterogeneous properties on resting-state dynamics.

**Fig. 4.**
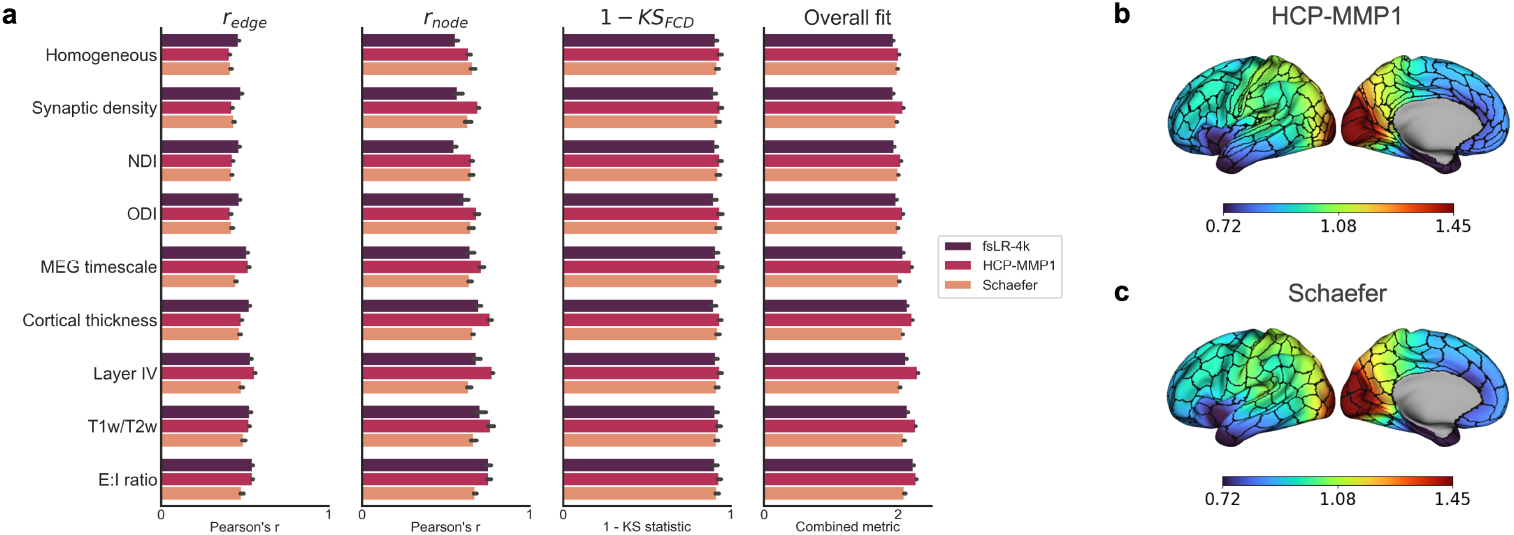
Impact of parcellation choice on model performance. **a**, Comparison of model fits when using vertex-level resolution (fsLR-4k) versus two popular atlas-based parcellations: HCP-MMP1 and Schaefer with 180 and 150 regions in one hemisphere, respectively. Groups are ordered from top to bottom by the mean overall fit of that map, averaged across the three models (fsLR-4k, HCP-MMP1, and Schaefer). Within each group, the three bars show the model-specific fits. **b**, HCP-MMP1 parcellation overlaid on the E:I ratio map. **c**, Schaefer parcellation overlaid on the E:I ratio map.

### The effects of spatial heterogeneity on wave dynamics are conserved across non-human primates

Having established the utility of accounting for cortical heterogeneity in humans, we now examine whether the same principles generalise to non-human primates. Specifically, we apply our heterogeneous wave modelling approach to two species—the rhesus macaque (*Macaca mulatta*) and the common marmoset (*Callithrix jacchus*)—which have been extensively analysed in previous NMM-based studies^73-75^ but not in the context of NFT. This analysis thus represents an important test of the generality of our wave model, given that the macaque and marmoset cortical surfaces have very different geometries compared to the human cortex; the marmoset cortex is essentially lissencephalic, contrasting with the highly folded human cortex, while the macaque cortex is an intermediate between these two extremes.

In the macaque, we evaluate 19 different heterogeneity maps, spanning structural (cortical thickness)^76^, myeloarchitectonic (T1w/T2w ratio)^76^, and chemoarchitectonic features (estimates of the densities of 16 different neurotransmitter receptors derived from receptor autoradiography)^48^. In the marmoset, we consider three heterogeneity maps: T1w/T2w ratio, cortical thickness, and cell density derived from Nissl staining, since brain-wide autoradiography data are not available for this species^78,79^. For both species, we model their cortical dynamics on their left cortical surfaces with 4002 vertices, matching the vertex count of the human mesh (see Methods).

The correlation structure of the macaque heterogeneity maps reveals three major clusters (Fig. 5a). One cluster groups intracortical myelin content (T1w/T2w) with receptors associated with excitatory neurotransmission (AMPA, Kainate, avg. excitatory, E:I ratio). This combination likely reflects a shared gradient aligned with the cortical hierarchy in this animal, given prior work indicating that myelin and excitatory markers co-vary with regional hierarchical ranks^22^. A second cluster is dominated by receptors associated with inhibitory GABAergic neurotransmission (GABA-A/BZ, GABA-B) and muscarinic receptors (M_1_, M_3_). A third cluster groups receptors associated with modulatory neurotransmitter systems (5-HT2, *α*_2_) with GABA-A. Other maps, such as cortical thickness and M_2_ receptor density, are weakly correlated and do not fall into any group.

**Fig. 5.**
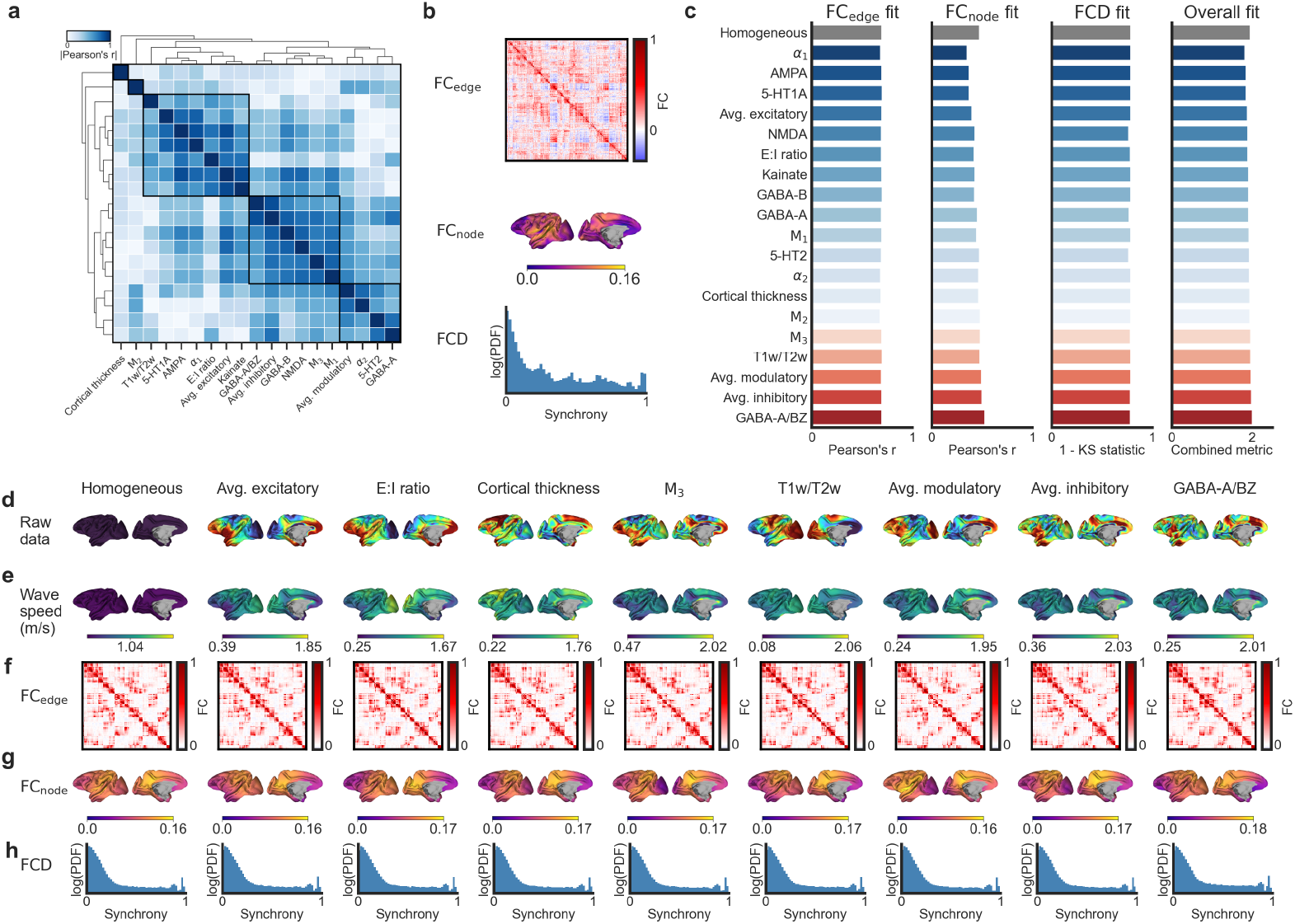
Generalisation of the heterogeneous wave model to the macaque cortex. **a**, Inter-map similarity (absolute Pearson’s correlation) across cortical heterogeneity maps. Hierarchical clustering highlights three dominant clusters of strongly related maps. **b**, Empirical vertex-level target properties used for model fitting and evaluation: (top) group-averaged edge-level functional coupling (FC) matrix (*FC*_*edge*_), (middle) vertex-wise node-level FC (*FC*_*node*_), and (bottom) distribution of functional connectivity dynamics (FCD; log-scaled y-axis). **c**, Fits for each model. The homogeneous model is shown for reference (grey), followed by the heterogeneous models ordered from lowest to highest overall fit (blue: lower overall fit than homogeneous; red: higher overall fit than homogeneous). The FCD fit is shown as 1 - KS statistic since lower KS values indicate better fit. The overall fit is calculated as ℒ = *r*_*edge*_ + *r*_*node*_ + (1 − *KS*_*FCD*_). **d-h**, Model inputs and evaluation outputs for the homogeneous model, followed by the four worst- and best performing heterogeneous models based on the overall fit (see Supp Fig. S4 for all other heterogeneous models). **d**, Raw heterogeneity maps displayed on the macaque cortical surface. **e**, Resulting spatial maps of wave speeds derived by using the optimised *α* parameter (see Methods). **f-h**, Model-derived (f) edge-level FC matrices, (g) node-level FC maps, and (h) FCD distributions (log-scaled y-axis) all defined at the vertex level. Panels d-h are arranged from left to right following the ordering in panel c.

We fit each model to the group-average edge-level FC and node-level FC, and the FCD calculated across the concatenated timeseries of awake resting-state fMRI signals recorded from each of the 10 macaques (Fig. 5b; see Methods). Compared to the human findings, the differences in performance between the homogeneous and heterogeneous models are smaller. Indeed, just under half of the heterogeneous models perform better than the homogeneous model, which shows an edge-level FC correlation with empirical data of *r* = 0.700 and a node-level FC correlation of *r* = 0.482. The optimised *α* values are generally close to zero (range -0.5 to +0.5), indicating that only subtle deviations from the homogeneous model are needed to improve model fits in the macaque. All models converge on a similar axonal length scale (*r*_*s*_≈ 9 mm) that is notably smaller than the human estimate, consistent with the shorter propagation ranges required in smaller brains.

The model parameterised by the inhibitory GABA-A/BZ receptor has the best overall fit, with the largest improvement relative to the homogeneous observed for node-level FC fit (Fig. 5c,f, and g; *r*_*edge*_ = 0.702 and *r*_*node*_ = 0.534) although again, the improvements are smaller (0.1% for *r*_*edge*_ and 10.8% for *r*_*node*_ compared to 26.0% for *r*_*edge*_ and 34.7% for *r*_*node*_, respectively, for the contrast between the human homogeneous and E:I balance model). The performances of the average inhibitory (*r*_*edge*_ = 0.702, *r*_*node*_ = 0.505), average modulatory (*r*_*edge*_ = 0.696, *r*_*node*_ = 0.500), and T1w/T2w models (*r*_*edge*_ = 0.707, *r*_*node*_ = 0.484) are very similar. The stronger performance of the T1w/T2w model aligns with the human findings but the gains are also smaller than observed in humans (e.g., 0.7% and 0.4% improvement for *r*_*edge*_ and *r*_*node*_ in macaques compared to 22.4% and 25.8% in humans, respectively). Notably, the E:I balance model does not surpass the performance of the homogeneous model in macaques (Fig. 5c), which may reflect either the different measures and/or receptors used to measure E:I balance in the two species. Notably, the best performing GABA-A/BZ map has an absolute Pearson’s correlation of 0.09 with the T1w/T2w ratio, indicating that these high-performing models capture distinct spatial patterns of heterogeneity and may represent multiple complementary constraints on cortical dynamics. The modelling revealed that regions with high GABA-A/BZ receptor density are associated with slower wave propagation (*α* < 0), whereas regions with high T1w/T2w ratio and cortical thickness are associated with faster propagation (*α* > 0), consistent with findings in humans (Fig. 5d,e). Consistent with the observations in humans, the addition of spatial heterogeneity in macaques produces minimal changes in FCD (Fig. 5c,h).

In the marmoset, we observe low spatial correlations between all heterogeneity maps, with the highest similarity between cell density and T1w/T2w (*r* = 0.38; Fig. 6a). Each of these maps thus captures distinct aspects of cortical heterogeneity. Comparison of the models reveals that incorporating intracortical myelin-related heterogeneity (T1w/T2w ratio) provides the largest improvement in model fit relative to the homogeneous baseline (Fig. 6c). We fit each model to the group-average edge-level FC and node-level FC, and the FCD calculated across the concatenated timeseries of awake resting-state fMRI signals recorded from each of the 39 marmosets (Fig. 6b; see Methods). Fitted *α* values indicate that regions with high T1w/T2w ratio and cortical thickness are associated with faster propagation (*α* > 0) whereas regions with high cell density are associated with slower wave propagation (*α* < 0). All models converge on a smaller axonal length scale (*r*_*s*_≈ 1 mm) than the other two species which is again proportional to the smaller size of the animal.

**Fig. 6.**
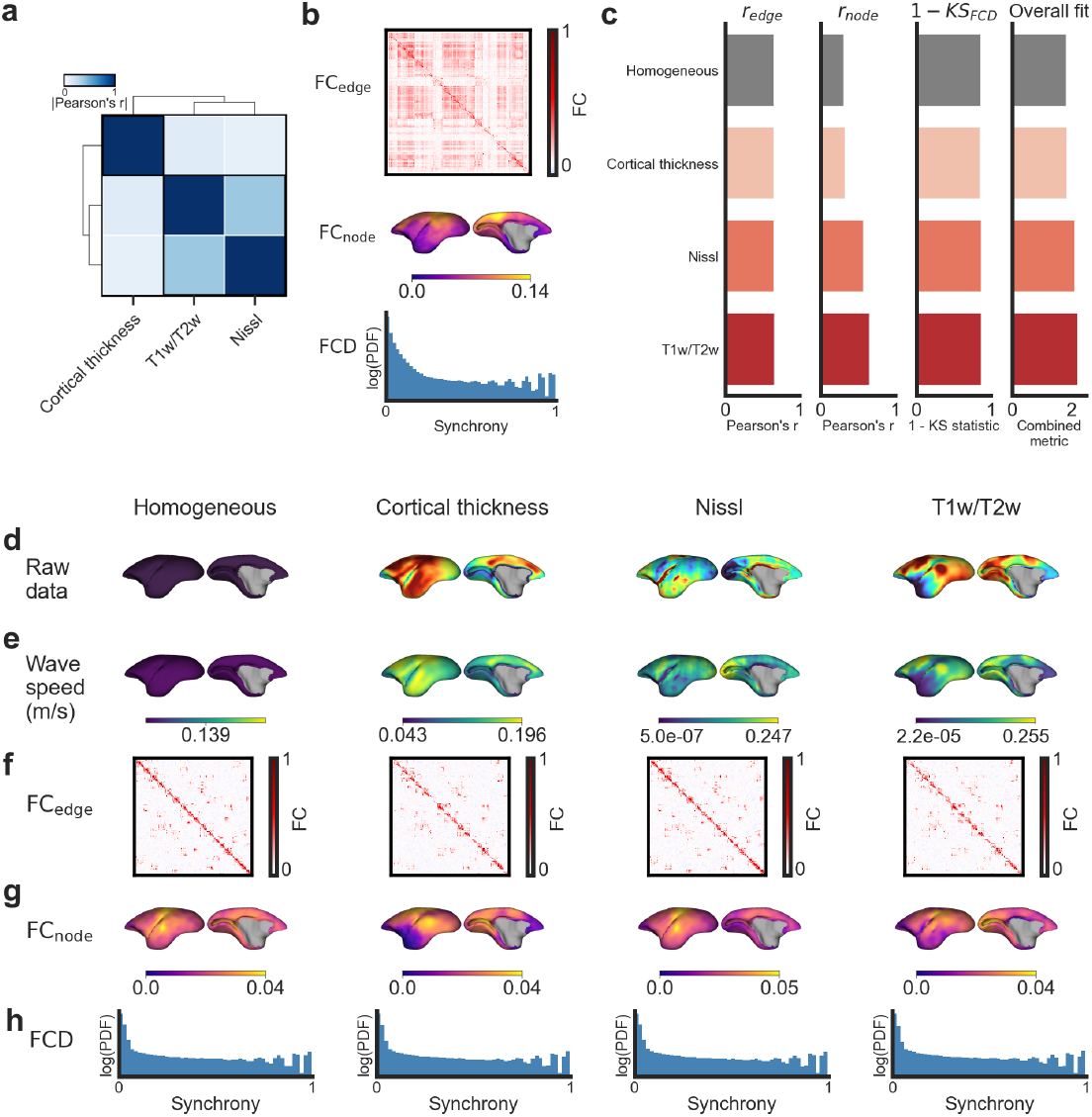
Generalisation of the heterogeneous wave model to the marmoset cortex. a, Inter-map similarity (absolute Pearson’s correlation) across cortical heterogeneity maps. **b**, Empirical vertex-level target properties used for model fitting and evaluation: (top) group-averaged edge-level functional coupling (FC) matrix (*FC*_*edge*_), (middle) vertex-wise node-level FC (*FC*_*node*_), and (bottom) distribution of functional connectivity dynamics (FCD; log-scaled y-axis). **c**, Fits for each model. The homogeneous model is shown for reference (grey), followed by the heterogeneous models ordered from lowest to highest overall fit. The FCD fit is shown as 1 - KS statistic since lower KS values indicate a better fit. The overall fit is calculated as ℒ = *r*_*edge*_ + *r*_*node*_ + (1 − *KS*_*FCD*_). **d-h**, Model inputs and evaluation outputs for the homogeneous and heterogeneous models. **d**, Raw heterogeneity maps displayed on the marmoset cortical surface. **e**, Resulting spatial maps of wave speeds derived by using the optimised *α* parameter (see Methods). **f-h**, Model-derived (f) edge-level FC matrices, (g) node-level FC maps, and (h) FCD distributions (log-scaled y-axis) all defined at the vertex level. Panels d-h are arranged from left to right following the ordering in panel c.

Across all models for marmoset, we again observe that the largest increases in model fit arise from an improvement in capturing edge-level FC (Fig. 6c,f) and node-level FC (Fig. 6c,g) with minimal variation in FCD fit (Fig. 6c,h). The largest gain is observed for node FC, where incorporating the T1w/T2w maps produces a 107.7% improvement relative to the homogeneous model. These findings indicate that even in a lissencephalic animal with relatively simple cortical geometry, local myeloarchitectonic variation exerts a conserved role in shaping macroscopic wave dynamics.

## Discussion

We present an NFT-based wave model that accommodates regional heterogeneities in the cortical medium. Unlike traditional NMMs, which discretise the cortex into several hundred regions, our approach treats the cortical surface as a continuous sheet, allowing us to model how fine-grained spatial variations in local cortical properties influence macroscopic wave dynamics. Our model affords two key capabilities: (i) high resolution simulations at the vertex level (>4000 points), which avoid issues related to predefined parcellations and can capture subtle changes in cortical organization; and (ii) dependence only on cortical geometry and a heterogeneity map, which greatly facilitates comparative modeling, providing a biologically grounded and computationally efficient way to investigate how local microstructural and molecular properties shape large-scale neural dynamics in the brain across diverse species. Our approach thus offers a general framework for identifying conserved and divergent organisational principles shaping large-scale cortical dynamics. Indeed, the strong performance of our high-resolution wave model points to the important influence of wave dynamics in shaping macroscale activity such as those captured in fMRI^32,46^. Indeed, wave dynamics also naturally emerge in NMMs, despite the discretisation imposed by the parcellation procedures on which they rely^33^, suggesting that waves are a fundamental element of macroscale cortical activity.

In all species, incorporating spatial heterogeneity into the wave model improved correspondence with empirical FC compared to homogeneous benchmark models. The largest gains occurred for maps parametrising heterogeneity with either the T1w/T2w ratio or a measure of local excitability and/or inhibitory capacity. In humans, the E:I ratio and T1w/T2w maps represent related, yet distinct, axes of hierarchical variation in the cortex, given their moderate correlation (*r* = 0.57 in human; *r* = 0.61 in macaque). The success of the E:I ratio model is consistent with past work showing that variations in transcriptional markers of E:I balance^23,25^ and/or local recurrent excitation strength^24^ generally improve the performance of NMMs. Regions with a higher E:I ratio indicate a relative predominance of excitatory over inhibitory transcriptional markers. Although this measure does not distinguish whether these markers are expressed on excitatory or inhibitory neurons, prior work has associated such E:I gradients with stronger local recurrent excitation and faster signal propagation^32^ .

The human E:I balance map was generated by aggregating transcriptional data across 8 genes coding for excitatory receptor subunits and 18 genes coding for inhibitory receptor subunits, whereas the available macaque data included only 3 excitatory and 3 inhibitory receptors measured using autoradiographic receptor-density estimates (see Methods). These two methods are fundamentally different since transcriptional markers provide an indirect proxy for protein abundance^80^, whereas receptor-density measurements in the macaque directly reflect protein levels but cover a more limited set of receptors compared to the human model. It is therefore unclear whether the discrepant findings in our human and macaque analyses reflect true differences in the effects of local E:I balance on wave dynamics between the two species or arise from differences in the methods used to estimate local E:I balance.

The T1w/T2w model also performed well, which is expected given the established link between the T1w/T2w ratio and intracortical myelin^21,81,82^ and prior evidence that this map serves as a proxy for cortical hierarchical rank along a unimodal-to-transmodal axis, which explains many spatially varying properties in the human cortex^22^. Our findings are also consistent with past work reporting substantial improvements in edge-level FC fit and node-level FC fit when incorporating T1w/T2w-based heterogeneity into NMMs^20,83^. Indeed, learning regional variations in the recurrent connection strength of an NMM yielded a spatial pattern that matched the same cortical hierarchy and which is positively correlated with empirical T1w/T2w estimates^24^. Accordingly, association areas exhibit more prolonged activity timescales and greater integrative capacity, whereas sensory regions support faster, more stimulus-driven responses^8,13,84-87^. Our observation that maps parameterised with T1w/T2w maps also improved fits to empirical fMRI recordings in macaque and marmoset indicates that the hierarchical gradient of myelination represents a highly conserved axis of organisation that constrains wave dynamics across primate brains with diverse cortical geometries and phylogenetic distances spanning millions of years.

In general, we found that the performance gains for heterogeneous models were greater in humans compared to macaques and marmosets. This result may reflect observations that cortical processing hierarchies, and thus, local variations in cellular, molecular, and physiological architecture, are deeper in animals with larger brains as a result of their extended period of postnatal neurodevelopment^3,88,89^. Such an extension augments regional heterogeneities in cellular, molecular, and functional architecture and may have been selected to support ongoing myelination and plasticity^88^. This view is supported by our observation that, consistent with previous work in NMMs^20^, performance gains of heterogeneous compared to homogeneous models applied to human data were most pronounced in transmodal association cortices. These areas exhibit greater variability in microstructural and molecular properties^90-92^ and also show a protracted period of postnatal maturation^3,71^. In contrast, earlier-maturing sensory areas are relatively uniform, less heterogeneous, and show smaller improvements in our analysis, meaning that even a simple homogeneous model can approximate dynamics in these areas reasonably well. However, we caution that the empirical fMRI data in non-human species were derived from small samples (i.e., 10 macaques and 39 marmosets compared to 255 humans), which may yield noisier FC estimates and reduce sensitivity to the effects of heterogeneity in the macaque and marmoset.

A particular advantage of our approach is that it does not require an a priori parcellation of the cortex. This is important, given our finding that the results of atlas-based models will depend on the degree to which parcel borders align with the underlying heterogeneity map—a condition that is difficult to guarantee across atlases. High-resolution models can overcome this problem and are computationally feasible within our NFT wave model framework. We demonstrate that our vertex-level NFT model achieves similar or better fits at substantially higher spatial resolution (e.g., compare our fit statistics with those reported in refs.^20,24^). An additional advantage is that our approach does not depend on detailed structural connectivity estimates derived from diffusion MRI, which introduce additional complexity and potential sources of error^30,93-95^. The need to measure connectivity also limits the ability to compare across species, where comprehensive connectome measures may not always be available. The strong performance of our model further reinforces prior work indicating that a simple isotropic, distance-dependent connectivity kernel following the exponential distance rule is sufficient to capture many different aspects of large-scale functional dynamics mapped with fMRI^46^. More complex features of connectivity that do not conform to this kernel may drive more rapid dynamics that are not readily accessible with fMRI^70,96-98^.

We used FCD as one of our evaluation metrics because it is widely adopted in the literature; however, FCD primarily captures temporal statistics and does not explicitly account for spatiotemporal patterns of network interactions. Future work could incorporate metrics that better quantify time-resolved variations in the spatial patterns of FC dynamics^32^. Additionally, our model assumes an isotropic wave propagation across the cortical medium, whereas biological connectivity is locally anisotropic. Extending the model to include anisotropic propagation could provide a more realistic representation of cortical dynamics. Finally, our approach focuses exclusively on cortico-cortical interactions and does not account for subcortical connectivity, which plays an important role in shaping large-scale brain activity^99-101^. Recent work has begun to address this limitation by incorporating subcortical inputs into NFT models^47^, and similar extensions could enhance the generality and biological plausibility of our framework.

In conclusion, our general framework for modelling the effects of regional heterogeneity on cortical dynamics identifies a highly conserved role for myeloarchitecture and regional E:I balance in shaping cortical activity. Our work demonstrates the importance of moving beyond models that assume uniform properties across brain regions by incorporating biologically informed spatial heterogeneity. This approach enables more accurate predictions of functional connectivity, captures hierarchical gradients of cortical organisation, and generalises across species with vastly different brain geometries.

## Methods

For each species, analyses were restricted to the left hemisphere. The only model inputs were a 2D triangular mesh of the cortical surface derived from T1w MRI and a spatial heterogeneity map. Resting-state fMRI was used to evaluate model performance. The heterogeneity maps and resting-state fMRI data were projected onto the corresponding cortical surface template of each species.

### Human dataset

#### Cortical mesh and heterogeneity maps

We modelled cortical dynamics on the left hemisphere of the human cortical surface. To facilitate group-level analyses, we used the left-right symmetric fsaverage midthickness template developed by FreeSurfer and HCP^48,49^. This template provides a standardised cortical geometry derived from T1w MRI data, enabling consistent projection of data across subjects. For computational tractability, we used a downsampled surface with 4002 vertices within a hemisphere (fsLR-4k), obtained from the neuromaps repository^10^. We used the template cortical surface rather than subject-specific surfaces for simplicity and because our primary goal was to test the effects of spatial heterogeneity on wave dynamics. Many more group-averaged (compared to individual-level) maps are available in the public domain, allowing us to more comprehensively assess diverse aspects of cortical heterogeneity. Specifically, we used eight distinct group-average cortical heterogeneity maps, each capturing a different molecular, cellular, microstructural, anatomical, or physiological property hypothesised to influence local wave speed, as encoded by our model. Each of these maps was projected onto the fsLR-4k surface and used to generate a distinct heterogeneous model, allowing us to systematically investigate how different forms of spatial heterogeneity shape macroscopic wave dynamics.

The T1w/T2w ratio, cortical thickness, and MEG timescale maps were all obtained from the neuromaps repository^10^, which were originally sourced from the HCP dataset^56^. The T1w/T2w ratio was derived from T1w and T2w MRI and serves as a proxy for intracortical myelin content^21,81,82^ and is putatively related to axonal conduction velocity. Cortical thickness was derived from T1w MRI and reflects regional microstructural organisation, including neuronal density and packing. The MEG-derived intrinsic timescale map provides a measure of the regional temporal integration window, reflecting dynamic preferences and how long a region retains information. Synaptic density was estimated using high-resolution [^11^C]UCB-J PET imaging of SV2A and obtained from prior published work^57^; this map approximates the regional capacity for local information transfer. Two maps were derived from gene expression data based on spatially dense cortical expression maps^58^ from the Allen Human Brain Atlas^102^: one representing layer IV gene expression, computed as the first principal component (PC) of genes expressed in cortical layer IV, and another representing the excitatory/inhibitory (E:I) ratio, defined as the first PC of marker genes for excitatory neurons divided by the first PC of marker genes for inhibitory neurons. These gene-derived maps approximate regional variations in thalamic input and local excitability, respectively. The layer IV gene list was obtained from ref.^58^ which was compiled from the union of layer-specific marker genes found in two comprehensive transcriptomic studies focused on layer-dependent gene expression in prefrontal cortical regions^103,104^. To derive the E:I ratio map, we first took the lists of marker genes for excitatory and inhibitory neurons from ref.^58^, which was compiled from multiple single-nucleus RNA sequencing datasets that each sampled various cortical lobes (occipital, temporal, frontal, cingulate and parietal; see ref.^58^ for details). These lists reflect the overall transcriptional profiles of excitatory and inhibitory neuronal subtypes, rather than representing genes solely coding for receptor subunits. To refine these lists, we filtered for genes annotated in the receptor complex by Gene Ontology (GO:0043235) in *Homo sapiens*, supported by direct experimental evidence manually curated from the literature^105,106^. We then computed the first PC of the maps in each refined gene set, scaled both PC maps to the range [1, 2], and expressed the E:I ratio as the excitatory PC divided by the inhibitory PC. The neurite density index (NDI) and orientation dispersion index (ODI) were obtained from the BALSA repository^107^, which were originally sourced from ref^59^. NDI indexes the proportion of voxel volume occupied by neurites and ODI indexes the heterogeneity of neurite fibre orientations, providing microstructural measures of local microstructural connectivity and anisotropy.

#### Resting-state fMRI

We used preprocessed, resting-state fMRI data from 255 unrelated healthy individuals (aged 22-35 years; 132 females and 123 males) from the HCP 1200 subject data release^64,108^, where imaging was performed using a 3 T scanner with 2.0 mm isotropic resolution, TR = 720 ms, and TE = 33 ms. The data were preprocessed according to the HCP minimal preprocessing pipeline^109^. For each participant, we used the left-to-right phase encoding time series from the first session (14.4 minutes; 1200 time frames), and downsampled it to the first 600 time frames after confirming that this did not affect model evaluation. This reduction substantially alleviated a major computational bottleneck in our framework: computing vertex-level FCD. The empirical targets for model evaluation (see Methods: Model fitting and optimisation) were then computed on a concatenated cohort-level time series comprising these 600 time frames from each of the 255 subjects (600 × 255 = 153,000 time frames).

### Macaque dataset

#### Cortical mesh and heterogeneity maps

The macaque cortical surface was represented using the Yerkes19 midthickness template^110^ downsampled to 4k for consistency with human data. Similarly to the human dataset, we used T1w/T2w ratio and cortical thickness maps, obtained from ref.^76^, to encode spatial heterogeneity. Additionally, we used 16 chemoarchitecture maps derived from autoradiographic imaging of neurotransmitter receptors for acetylcholine (M_1_, M_2_, and M_3_), GABA (GABA-A, GABA-A/BZ, and GABA-B), glutamate (AMPA, Kainate, and NMDA), noradrenaline (*α*_1_ and *α*_2_), and serotonin (5-HT1A, 5-HT2). We also used three summary maps representing the average density of receptors associated with (i) excitatory neurotransmission (glutamatergic receptors), (ii) inhibitory neurotransmission (GABAergic receptors), and (iii) modulatory neurotransmitter systems (serotonin, acetylcholine, and noradrenaline). Here, ‘modulatory’ refers to the neurotransmitter systems and not the excitatory or inhibitory actions of specific receptor subtypes. These receptor maps were included because neurotransmitter receptor density is thought to influence local excitability and synaptic dynamics and thus may modulate wave speed in the NFT wave model. The receptor density data come from ongoing, unpublished work, with estimates reconstructed using the processing pipeline described in ref.^77^. We also computed an E:I ratio map—conceptually a glutamatergic:GABAergic ratio—by dividing the average excitatory map by the average inhibitory map, after scaling both maps to the range [1, 2] to avoid division by zero. This resulted in a total of 19 heterogeneity maps for the macaque.

#### Resting-state fMRI

We used preprocessed resting-state fMRI data from 10 awake macaques^111^, where the imaging was performed using a 4.7 T scanner, TR = 2600 ms, TE = 7 ms, and two scan sessions each with a total duration of 10.8 minutes (250 time frames). To yield a similar number of time frames in the marmoset dataset discussed below, we concatenated both sessions which resulted in a total duration of 21.6 minutes (500 time frames). The fMRI preprocessing included slice timing correction, motion correction, nuisance regression, band-pass filtering of 0.01-0.1 Hz, and surface smoothing using a 3-mm FWHM Gaussian kernel.

### Marmoset dataset

#### Cortical mesh and heterogeneity maps

The marmoset cortical surface was represented using the Marmoset Brain Mapping v3 (MBMv3) midthickness template^78^ downsampled to 4k for consistency with human and macaque data. The T1w/T2w ratio and cortical thickness maps were also obtained from the MBMv3, while the cell density map derived from Nissl staining was obtained from refs.^11,79^. We used the Nissl-stained map to test the effects that cell density would have on local excitability and signal propagation in the NFT wave model.

#### Resting-state fMRI

We used preprocessed resting-state fMRI from 39 awake marmosets across two independent sites: the Institute of Neuroscience (ION, China) and the National Institutes of Health (NIH, USA). At the ION site, imaging was performed on a 9.4 T scanner with 0.5 mm isotropic resolution, TR = 2000 ms, TE = 18 ms, and a total duration of 17.1 minutes (512 time frames). At the NIH site, data were acquired on a 7 T scanner with 0.5 mm isotropic resolution, TR = 2000 ms, TE = 22.2 ms, and a total duration of 17.1 minutes (512 time frames). Full acquisition details are available in ref.^112^. The fMRI preprocessing included slice timing correction, motion correction, nuisance regression, band-pass filtering of 0.01-0.1 Hz, and surface smoothing using a 2-mm FWHM Gaussian kernel.

#### The spatially heterogeneous wave model

To incorporate spatial heterogeneity into the NFT wave model, we used the generalised form of the damped wave equation (Eq. 1). This equation models the spatiotemporal evolution of mean-field population activity across the cortical surface, treated as a spatially continuous medium. Neural activity is represented as waves that propagate through this medium, shaped by connectivity and cortical geometry. The external input term 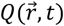 drives the system, representing external stimuli or intrinsic fluctuations.

Our analysis focused on the spatial component of the wave dynamics modelled by Eq. 1, which is determined by the LBO. The LBO characterises how functions vary across a curved surface by incorporating the local geometry of the mesh, quantifying relationships between neighbouring vertices in terms of their spatial distances and surface curvature. Its continuous definition is given by the divergence of the surface gradient, which in local coordinates takes the form

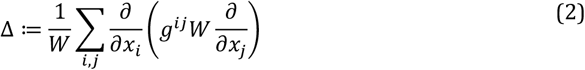

where *x*_*i*_, *x*_*j*_ are the local coordinates, *g*^*ij*^ is the inverse of the inner product metric tensor 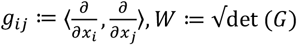, det denotes the determinant, and *G* ≔ (*g*_*ij*_ ). In practice, the continuous LBO (Eq. 2) is discretised on the cortical mesh using a finite element method.

Importantly, Eq. 1 generalises the NFT wave equation developed previously^36,46^ by introducing a position-dependent diffusion tensor, 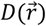, that allows us to embed spatial heterogeneity into the LBO. This formulation is based on the anisotropic LBO originally developed for shape analysis^113^. We adapt this approach by restricting the diffusion tensor to be isotropic (equal values on the diagonals) and allowing the tensor to vary across the cortical surface, thus deriving a spatially heterogeneous LBO, Δ_*D*_. This modification has a direct impact on the wave dynamics. The baseline wave speed is defined as *γ*_*s*_*r*_*s*_, which has units in length/time. This term is rescaled by the diffusion tensor, so that the local wave speed is equal to 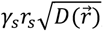, giving each cortical location a distinct wave speed based on the corresponding value of the heterogeneity map (e.g., T1w/T2w).

The diffusion tensor was defined as an isotropic, diagonal matrix:

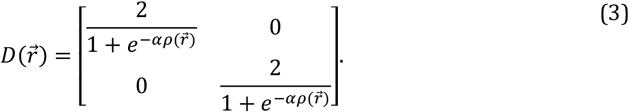

Each diagonal entry was scaled using a sigmoid transform of the heterogeneity map, 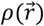, which was z-scored to ensure comparability across heterogeneity maps of different scales. The sigmoid transform maps values of 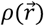 onto the interval (0, 2), which ensures that the diffusion tensor is positive-definite, thus yielding strictly positive diffusion values and real-valued wave speeds. This sigmoid transformation also truncates extreme values that could destabilise the computation. To tune the sensitivity of the wave speed to the underlying heterogeneity, we introduced a scaling parameter, *α*, which controls the steepness of the sigmoid curve. A large *α* makes the mapping highly nonlinear, thus accentuating differences in 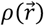, whereas a small *α* yields a nearly linear mapping, thus weakening the effect of the heterogeneity. When 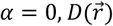 reduces to the identity matrix, and the heterogeneous LBO, Δ_*D*_, collapses to the homogeneous LBO, Δ, yielding homogeneous wave propagation. We also allowed *α* to take negative values, which inverts the relationship between the heterogeneity map and wave speed. This allows the model to easily account for cortical properties where higher values result in slower wave speeds and lower values result in faster wave speeds.

#### Simulating resting-state neural dynamics

Following refs.^46,61-63^, we simulated spontaneous or so-called “resting state” neural activity by solving Eq. 1 with a Gaussian white noise external input to mimic unstructured stochastic fluctuations with no biases to a particular frequency. These inputs could represent endogenous and/or exogenous stimuli or the activity of other brain structures that are not explicitly modelled in the NFT wave model. We fixed *γ*_*s*_to 0.116 *ms*^−1^ as estimated from physiological measurements^114^. This choice was justified by secondary analyses indicating that variations of this parameter exert a weak impact on the resulting dynamics. This left *r*_*s*_and *α* as the only free parameters fitted to the data.

To efficiently solve Eq. 1 on the cortical surface, we exploited an eigenmode expansion, following previous work^46^. Specifically, since the white Gaussian input, 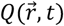, is linear we expressed it and the solution, 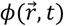, as linear combinations of a basis set, 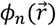:

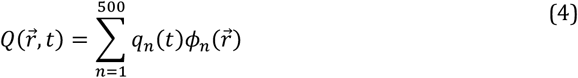

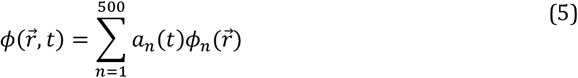

where 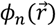 are the eigenmodes of the cortical mesh, which satisfy

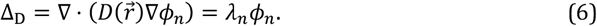

Note that we used the first 500 eigenmodes, consistent with previous work^46^, after validating that varying the number of modes did not significantly change the results of the study. Because we used a generalised LBO with a spatially varying diffusion tensor, the resulting eigenmodes were spatially heterogeneous and differed slightly from the typical geometric eigenmodes obtained using a spatially homogeneous diffusion tensor. We illustrate these differences in Fig. S5.

Substituting Eqs. 4 and 5 into Eq. 1 reduced the full spatiotemporal partial differential equation to a set of decoupled ordinary differential equations for the mode amplitudes:

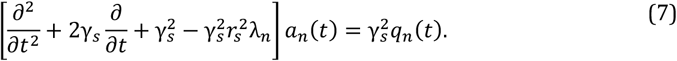

Here *q*_*n*_(*t*) is the projection of the input 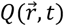 onto the *n*-th eigenmode. Each mode therefore evolved independently as a damped oscillator with a frequency that depends on the eigenvalue. Once solved, the modes and mode amplitudes were recombined via Eq. 5 to reconstruct the full neural activity. This approach substantially reduced computational cost while enabling stable, high-resolution simulations on the cortical mesh.

For the human models, the NFT wave model was simulated with a timestep of 90 ms. To match the 720 ms TR of the empirical fMRI data, model outputs were sampled every 8 simulation steps (720/90 = 8). The total number of simulation timepoints was therefore 600 × 8 = 4800, where 600 is the number of volumes in the empirical dataset. For the macaque and marmoset models, timesteps of 100 ms were used, and outputs were downsampled every 26 steps (macaque; 2600/100 ms) and every 20 steps (marmoset; 2000/100 ms) to match the empirical TRs. With 500 macaque TRs and 510 marmoset volumes, this corresponded to 500 × 20 = 10,000 and 510 × 26 = 13,260 simulation timepoints, respectively. In all species, the first 550 timepoints were discarded to allow the model to reach steady state.

We then converted the neural activity output of the NFT wave model to simulated blood oxygen-level dependent (BOLD) signals via the Balloon-Windkessel hemodynamic model^50,51^ for direct comparison with empirical fMRI measures:

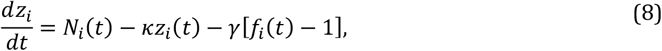

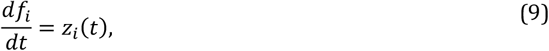

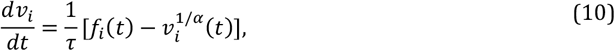

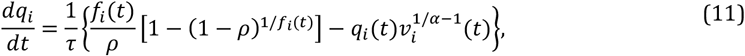

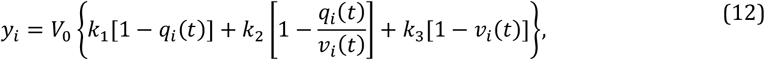

where *z*_*i*_, *f*_*i*_, *v*_*i*_, *q*_*i*_, and *y*_*i*_ are the vasodilatory signal, blood inflow, blood volume, deoxyhaemoglobin content, and BOLD signal variables, respectively. The model parameters were the same as the ones described in ref.^46^. We then z-scored the BOLD signal and fitted it to the empirical resting-state BOLD-fMRI data.

#### Model fitting and optimisation

We optimised model parameters *r*_*s*_and *α* to fit two static FC properties—static pairwise FC (edge-level FC) and static node-level average FC (node-level FC)—and one dynamic FC property, called functional connectivity dynamics (FCD). Edge-level FC fit, *r*_*edge*_, was calculated by computing Pearson’s correlation between the upper-triangular elements of the model and empirical FC matrices. Node-level FC fit, *r*_*node*_, was calculated by computing Pearson’s correlation between the mean FC strength for each cortical vertex between the model and empirical data. For both metrics, higher correlation values reflected better agreement between model and data.

To capture the temporal evolution of connectivity, we employed the FCD metric, *KS*_*FCD*_, as used in previous work^46^. The vertex-level time series were first band-pass filtered between 0.04 and 0.07 Hz using a second-order Butterworth filter where the range was chosen based on prior work^115^ and its relevance for functional brain dynamics. The analytic signal of each time series was then obtained via the Hilbert transform, yielding a complex-valued representation. From this representation, we computed the instantaneous phase for each vertex and the pairwise synchrony between regions at each time point as the cosine of the phase difference. The global synchrony at two time points was then defined as the average similarity of pairwise synchrony across all regions, forming a symmetric time × time matrix. This matrix captures the temporal structure of network-level interactions. Finally, we compared the distributions of FCD values from the model and empirical data by concatenating the upper-triangular elements across subjects or model realisations and quantified their similarity using the Kolmogorov-Smirnov (KS) statistic. Lower KS values indicate better agreement between model and data.

We defined a single objective function as:

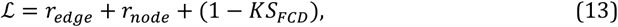

This function was maximised such that higher values indicated a better overall fit. To avoid over-fitting and allow fair comparison between the 2-parameter (*r*_*s*_and *α*) heterogeneous models and the 1-parameter (*r*_*s*_) homogeneous model, we used 5-fold cross-validation. In each fold, the free parameters were estimated using 80% of the subjects and the model fit was tested on the remaining 20%. The evaluation metrics reported for each model were calculated as the average test score across the 5 folds. For the macaque and marmoset analyses, where we only had a small number of individuals, we fitted the model directly to the data without cross-validation. We performed brute-force optimisation over a grid of parameter values. An initial coarse search was conducted over a broad parameter range to ensure adequate coverage of the parameter space (see Fig. S2). Based on this preliminary exploration, the final cross-validation analysis was performed over a restricted range with *α* ∈ [−3,3] in increments of 0.5 and *r*_*s*_∈ [15,20] in increments of 1.0.

To ensure robustness of the optimisation, we ran each model 10 times and computed the evaluation outputs on the concatenated timeseries from each run. This number was determined empirically, as preliminary testing showed that the objective function plateaued after approximately 10 runs, indicating diminishing gains in accuracy from additional runs. Edge-level FC, node-level FC, and FCD were subsequently computed from the concatenated timeseries across all runs.

#### Statistical inference

To generate appropriate null maps for statistical inference while controlling for the influence of spatial autocorrelation, we employed the eigenstrapping method^72^. Compared to traditional null map generation methods, such as the Spin test^116^ and BrainSMASH^117^, eigenstrapping explores a deeper null space, offering better control over false positives and better preservation of spatial autocorrelation. The methodology involved decomposing a heterogeneity map (e.g., T1w/T2w) using a spectral decomposition based on geometric eigenmodes derived from the homogeneous LBO. These eigenmodes, which represent spatial patterns at different scales, were partitioned into groups of approximately similar wavelengths. The geometric eigenmodes were then mapped to an equivalent spherical representation by normalising their eigenvalues and randomly rotating them within their respective groups using a random rotation matrix. Finally, surrogate maps were reconstructed from these rotated eigenmodes using the original decomposition coefficients, creating null maps that preserve the original spatial autocorrelation structure. For each heterogeneous model, we generated 500 null heterogeneity maps and ran the corresponding models through the same 5-fold cross-validation procedure described above. We constructed null distributions of the difference in model fit relative to the homogeneous model by computing, for each null realisation, the difference in fit between the null model and the homogeneous model. Significance was quantified as the fraction of null samples whose difference in fit equalled or exceeded the observed difference obtained with the empirical heterogeneity map (Fig. S3).

#### Network analysis

To assess the specificity of improved FC fits in certain areas of the human cortex, we compared the best performing heterogeneous model (E:I ratio) and the homogeneous baseline using the optimised FC matrices from the cross-validation procedure (see Simulating resting-state neural dynamics). Parcels were assigned to one of eight canonical human resting-state networks (RSNs) based on the community detection solution reported by ref.^20^, which was derived from the correlation structure of fMRI data in the HCP dataset using the HCP-MMP1 parcellation^56^. This classification yields three sensory networks (auditory, visual, somatomotor) and five association networks (dorsal attention, frontoparietal, ventral attention, default mode, and cingulo-opercular).

To evaluate vertex-level correspondence between empirical and model-generated FC, we computed Pearson’s correlation coefficients between corresponding rows of the empirical and simulated FC matrices. For each vertex, this yielded a measure of how well the model captured its connectivity profile with the rest of the cortex. Vertices were then grouped by RSN assignment and network-level improvements were quantified by comparing the distribution of vertex-wise accuracy values between the homogeneous and E:I ratio-based heterogeneous models. Differences in mean accuracy across networks were assessed using independent two-sample t-tests.

## Data and code availability

All source data and code to generate the results of this study will be made openly available upon publication. All analyses were implemented in Python, where we made use of the BrainSpace^118^ and LaPy libraries^119,120^, alongside other standard scientific libraries. Computations were performed using the MASSIVE high performance computing infrastructure^121^.

## Acknowledgements

This work was supported by Monash eResearch capabilities, including the M3 MASSIVE high-performance computing facility (www.massive.org.au). V.B. and I.Z.P were supported by an Australian Government Research Training Program (RTP) Scholarship (doi.org/10.82133/C42F-K220). N.P.-G. was supported by the Helmholtz Association’s Initiative and Networking Fund through the Helmholtz International BigBrain Analytics and Learning Laboratory (HIBALL) under the Helmholtz International Lab grant agreement InterLabs-0015. J.C.P. was supported by the Australian National Health and Medical Research Council (2034000) and Monash FMNHS Early Career Research Excellence Program. A.F. was supported by the Australian Research Council (FL220100184), National Health and Medical Research Council (1197431), and Sylvia and Charles Viertel Charitable Foundation (2017042). We thank Peter Robinson for insightful discussions during the early stages of this work.

