## Supplementary Figures S1-S5 for "Regional heterogeneity shapes macroscopic wave dynamics of the human and non-human primate cortex"

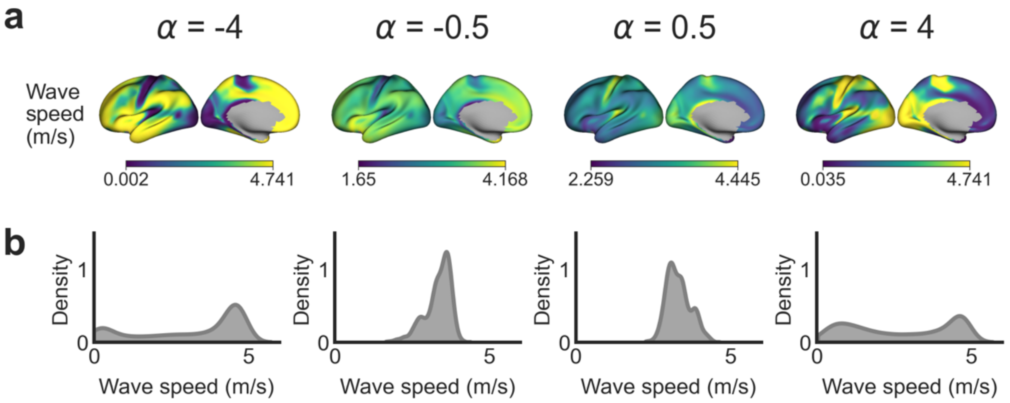


**Fig. S1 | Effects of alpha on heterogeneity map. a,** Resulting spatial maps of wave speeds using the human T1w/T2w model with varying $\alpha$ values. Note how $\alpha < 0$ inverts the map. **b,** Probability distribution function of the wave speed maps.

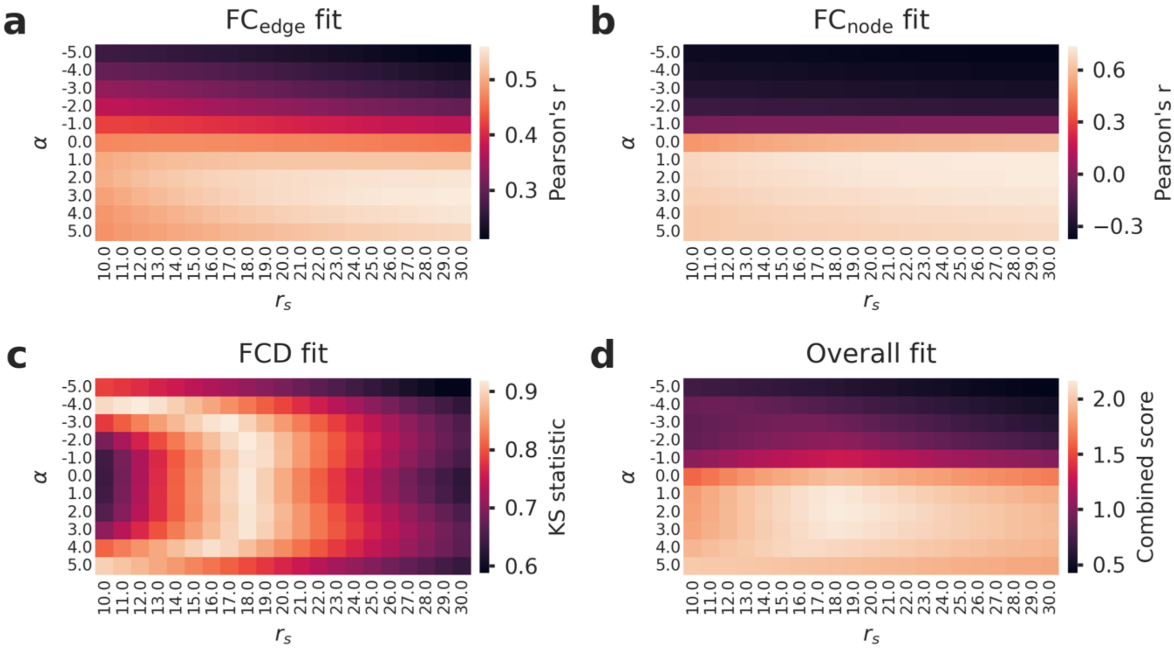


**Fig. S2 | Parameter landscapes for evaluation metrics. a-d,** Parameter landscapes for the (a) edge-level FC, (b) node-level FC, (c) FCD, and (d) overall fit when optimizing the human T1w/T2w model. Note that a finer $\alpha$-step of 0.5 was used for the final modelling results. This shows the broader landscape to ensure we correctly found a minimum.

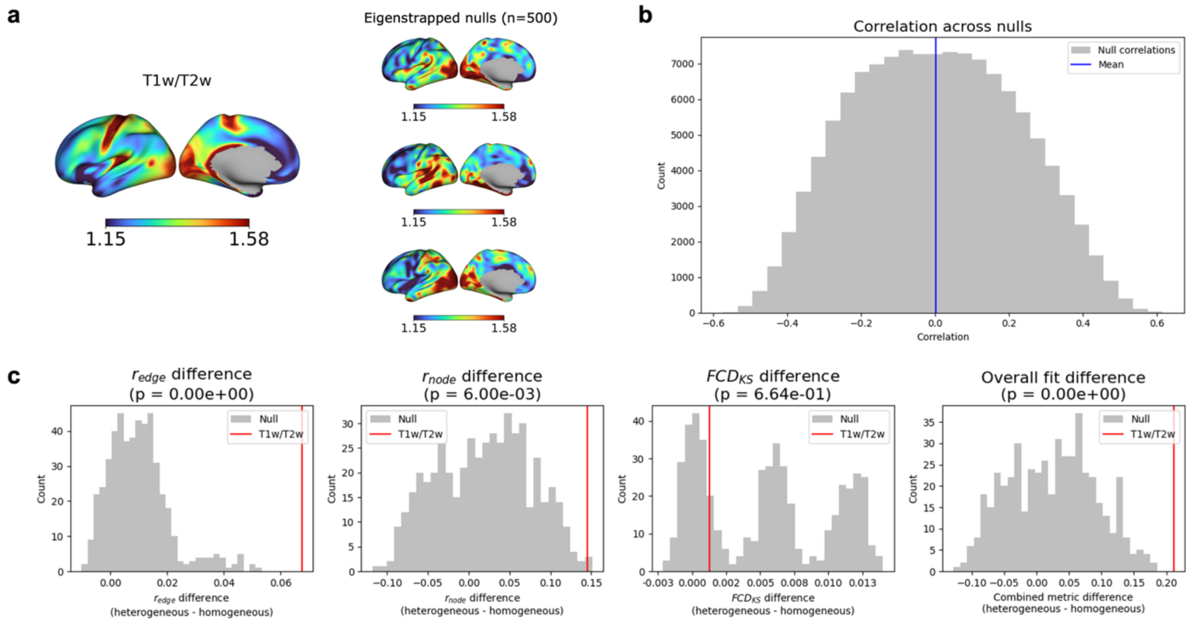


**Fig. S3 | Eigenstrapping method for statistical testing. a,** The input cortical map (T1w/T2w) to generate the nulls and 3 examples of the generate null maps. **b,** Histogram of spatial correlations (Pearson's r) between all pairs of nulls maps. **c,** Histogram of the fits for each metric for the null models. The T1w/T2w model significantly higher than the nulls.

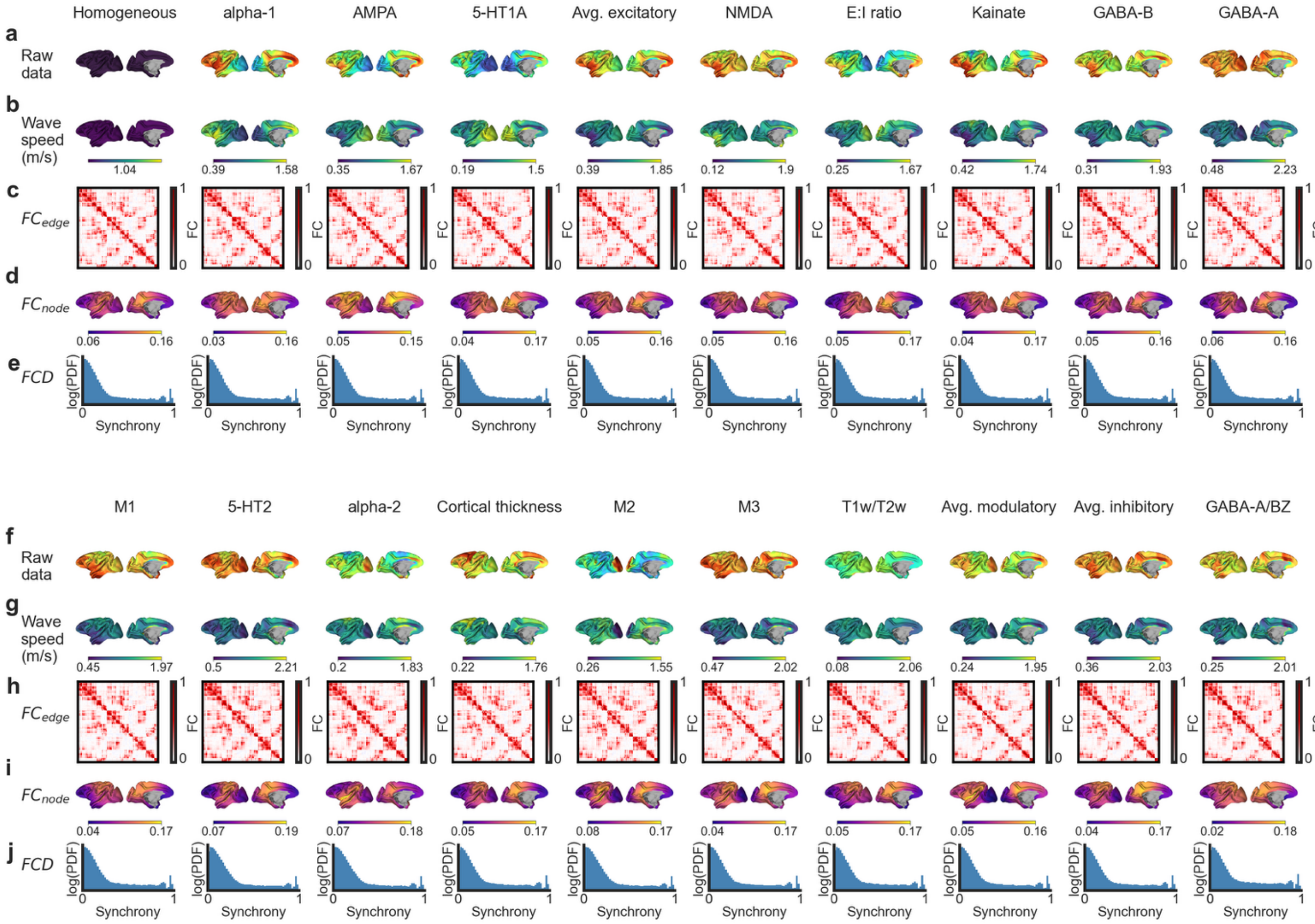


**Fig. S4 | Model inputs and evaluation outputs for all macaque models. a & f**, Raw heterogeneity maps displayed on the macaque cortical surface. **b & g**, Resulting spatial maps of wave speeds derived by using the optimized α parameter (see Methods). **c-e & h-j**, Model-derived (c and h) edge-level FC matrices, (d and i) node-level FC maps, and (e and j) FCD distributions (log-scaled y-axis) all defined at the vertex level.

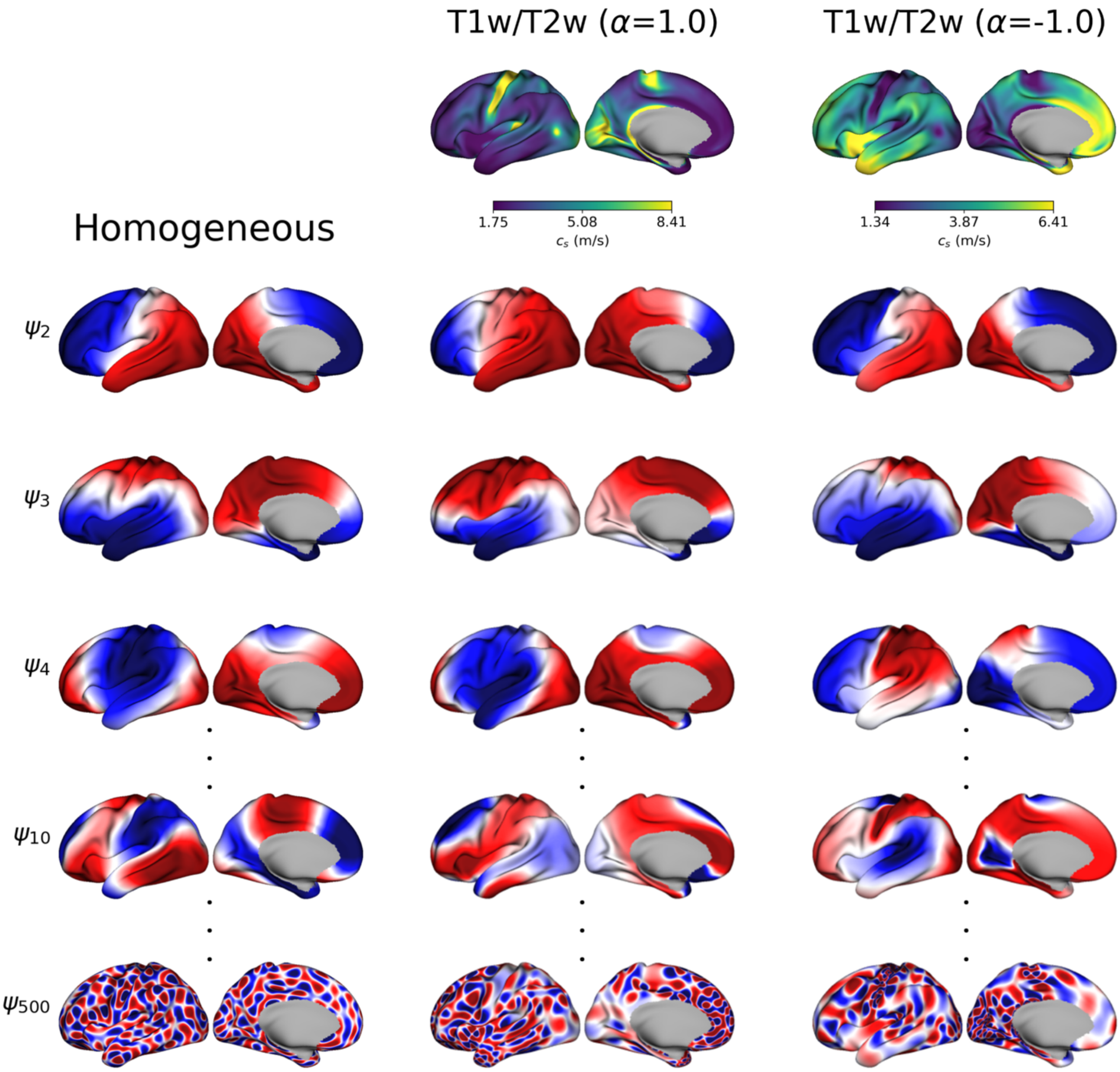


**Fig. S5 | Homogeneous and heterogeneous eigenmode basis sets.** The basis sets from left to right are homogeneous eigenmodes, heterogeneous eigenmodes parameterized by T1w/T2w with $\alpha = 1.0$, and heterogeneous eigenmodes parameterized by T1w/T2w with $\alpha = -1.0$. Negative-zero–positive values are coloured as blue–white–red. The wave speed maps for the heterogeneous eigenmodes are shown for reference.
